# eBiota: Designing microbial communities from large seed pools with desired function using rapid optimization and deep learning

**DOI:** 10.64898/2026.03.29.714676

**Authors:** Xiaoqing Jiang, Jiaheng Hou, Haoyu Zhang, Jinyuan Guo, Shaohua Gu, Doris Vandeputte, Yulin Liao, Qian Guo, Xinrun Yang, Yiyan Zhou, Peter X. Geng, Chunhui Wang, Mo Li, Alexandre Jousset, Xiaotao Shen, Zhong Wei, Huaiqiu Zhu

## Abstract

Designing microbial communities to generate target products is crucial for biotechnology, agriculture, and disease treatment. However, rationally designing such communities from large seed pools has become a major challenge, as the rapidly expanding number of complete microbial genomes greatly expands the search space and sharply increases the required screening time and computational cost. Here, we introduce eBiota, a platform for *ab initio* design of microbial communities from a pool of 21,514 strains to generate target products. eBiota not only identifies optimal strain combinations but also simulates community behaviors, including microbial interactions and relative abundances. eBiota integrates three modules: CoreBFS, a graph-based search algorithm that rapidly screens for bacteria with complete metabolic pathways related to the target product; ProdFBA, an extended flux balance analysis that identifies microbial consortia with maximal production efficiency; and DeepCooc, a deep learning model trained on 23,323 microbiome samples across various environments to infer co-occurrence patterns. We validated eBiota’s capabilities in microbial community design and production efficiency calculation using public microbiome datasets, ranging from single strains to six-member consortia. Further *in vitro* experiments involving 94 strains confirmed eBiota’s ability to identify species that inhibit pathogen growth and to accurately model the relative abundances within complex microbial communities. As an initial digital twin, eBiota provides a powerful platform for the rational design of functional microbial communities, offering new opportunities for metabolic engineering and synthetic biology.

## Introduction

Designing microbial factories to perform desired functions has long been a central pursuit in various fields, including energy biotechnology, sustainable agriculture, microbial therapeutics, and drug manufacturing^1–3^. The primary goal is to improve the production or degradation efficiency of target metabolites^4^. Although metabolic engineering and synthetic biology have led to great progress in engineering individual microbes^5^, single-strain systems often face intrinsic metabolic bottlenecks, by-product accumulation, and physiological constraints^6^. Microbial consortia offer a transformative alternative, allowing more efficient utilization of complex substrates, reduced genetic burden, and mitigation of by-product accumulation^7^. Meanwhile, rapid advances in high-throughput sequencing have greatly expanded our understanding of microbial diversity, vastly increasing the pool of candidate taxa and opening new possibilities for constructing functional microbial communities^8^.

Genome-scale metabolic models (GEMs) combined with flux balance analysis (FBA), as implemented in tools such as MICOM^9^, OptCom^10^, SteadyCom^11^, and Microbiome Modeling Toolbox^12^, have revolutionized microbial consortium modeling by enabling accurate prediction of substrate utilization, metabolite production, interactions, relative abundances and growth rates^13–16^. However, most existing methods remain largely descriptive and do not support the rational design of microbial communities^7^. Only a few pioneering efforts have attempted community design, falling into two main categories: graph-based methods (e.g., CoMiDA^17^ and MultiPus^18^) and FBA-based ones (e.g., FLYCOP^19^). Graph-based methods focus on the topology of microbial metabolic networks, making them computationally efficient but less effective^20^, whereas FLYCOP incorporates metabolic fluxes but requires prior consortium configurations. Therefore, there is an urgent need for new approaches to microbial community design, particularly given the vast pool of microbial candidates. Here, we introduce eBiota, which leverages graph-based method, FBA-based modeling, and deep learning to design microbial communities with specific functions from a large seed pool of over 20,000 strains.

The first bottleneck major in rational community design is the “curse of dimensionality” arising from selecting strains from large seed pools. Because time and computational cost increase exponentially with pool size, current approaches are typically restricted to seed pools of only several, or at most a few hundred strains^18,19^. As a result, a vast number of microorganisms are excluded from consideration despite having complete genomes or proteomes, leading to missed opportunities to identify communities with superior yields or novel functional outputs. To address this, eBiota incorporates CoreBFS, a graph-based search algorithm that largely reduces computational time and cost, enabling large-scale screening of candidate taxa.

A second challenge is identifying community members that can optimally perform the desired function, namely maximizing production or degradation efficiency. This requires accurate modeling of microbial communities with maximal production or degradation flux as the objective. Although classical FBA is widely used to compute flux distributions that optimize a specified objective, typically growth rate, it suffers from solution redundancy^21,22^ because multiple flux states can yield the same outcome. Parsimonious FBA (pFBA) alleviates this by introducing a secondary objective that minimizes total fluxes^23,24^. While several extensions^9–11^ enhance community-level growth modeling, they fall short when the objective is to maximize production or degradation efficiency, especially in communities where members trade off growth for metabolite exchange^9,25^. Flux variability analysis (FVA)^26,27^ instead quantifies feasible flux ranges under optimal growth and provides bounds on production or degradation flux, but it therefore lacks deterministic predictions. Thus, we developed ProdFBA within eBiota, which integrates the strengths of pFBA and FVA through multi-step optimization to generate deterministic flux distributions that achieve optimal production or degradation efficiency alongside near-maximal growth.

The third challenge is ensuring growth compatibility and ecological stability within designed microbial communities^7,28,29^. Co-occurrence patterns have been recognized as key indicators of stable coexistence^30–32^, and statistical tools such as SPIEC-EASI^33^ and SparCC^34^ are widely used to infer these patterns from abundance data. The heuristic GEM-based method HiOrCo^30^ further identifies higher-order co-occurrence patterns in large datasets, capturing complex multi-species associations. However, these statistical approaches cannot predict potential relationships involving novel taxa. Existing machine-learning models^35,36^, which are typically trained on limited datasets comprising only a few hundred species and restricted to pairwise associations, also exhibit limited generalizability. Fortunately, the vast amount of microbiome data from natural environments provides a rich foundation for understanding microbial co-occurrence at scale. Building on these resources, we developed DeepCooc within eBiota, a deep learning model trained on 23,323 microbiome samples spanning more than 1,500 genera, capable of predicting both pairwise and higher-order co-occurrence patterns in microbial communities.

Here, we present eBiota (https://github.com/allons42/eBiota), a platform for the *ab initio* design and behavioral modeling of microbial communities from large bacterial seed pools. By integrating CoreBFS, ProdFBA, and DeepCooc, eBiota designs communities from a pool of 21,514 strains capable of utilizing 309 substrates and producing 247 products, a scale that remains computationally prohibitive for existing tools but was achieved by eBiota within weeks. We validated eBiota using both public datasets and *in vitro* experiments. eBiota showed good performance in designing microbial consortia of one to six members for hydrogen production, selecting pathogen-inhibiting strains from 94 candidates, and accurately predicting their relative abundances. In summary, eBiota establishes a digital twin of the microbiome that aims to mimic the entire microbial community and simulate its diverse behaviors, setting it apart from more limited standard machine learning models. eBiota is hoped to serve as a valuable platform for the rational design of microbial communities with desired functions, guided by co-occurrence patterns.

## Results

### eBiota: a one-stop integration of three high performance tools

The workflow of eBiota is shown in **Fig. 1**. The input is a target metabolite to be produced or degraded. eBiota integrates three building blocks, namely CoreBFS, ProdFBA, and DeepCooc. CoreBFS rapidly screens bacterial candidates with complete (sub)pathways related to the target metabolite. ProdFBA further identifies bacterial candidates with high production or degradation efficiency while maintaining enough growth rate. The selected candidates are subsequently assembled into microbial communities through intermediates. ProdFBA further filters these communities for high functional efficiency and growth rate, and DeepCooc predicts co-occurrence patterns among members. eBiota finally outputs detailed information on the designed microbial consortia, including production or degradation efficiency, yield, interactions, cross-feedings, relative abundance, substrates, products, culture media, and growth rate.

**Fig. 1.**
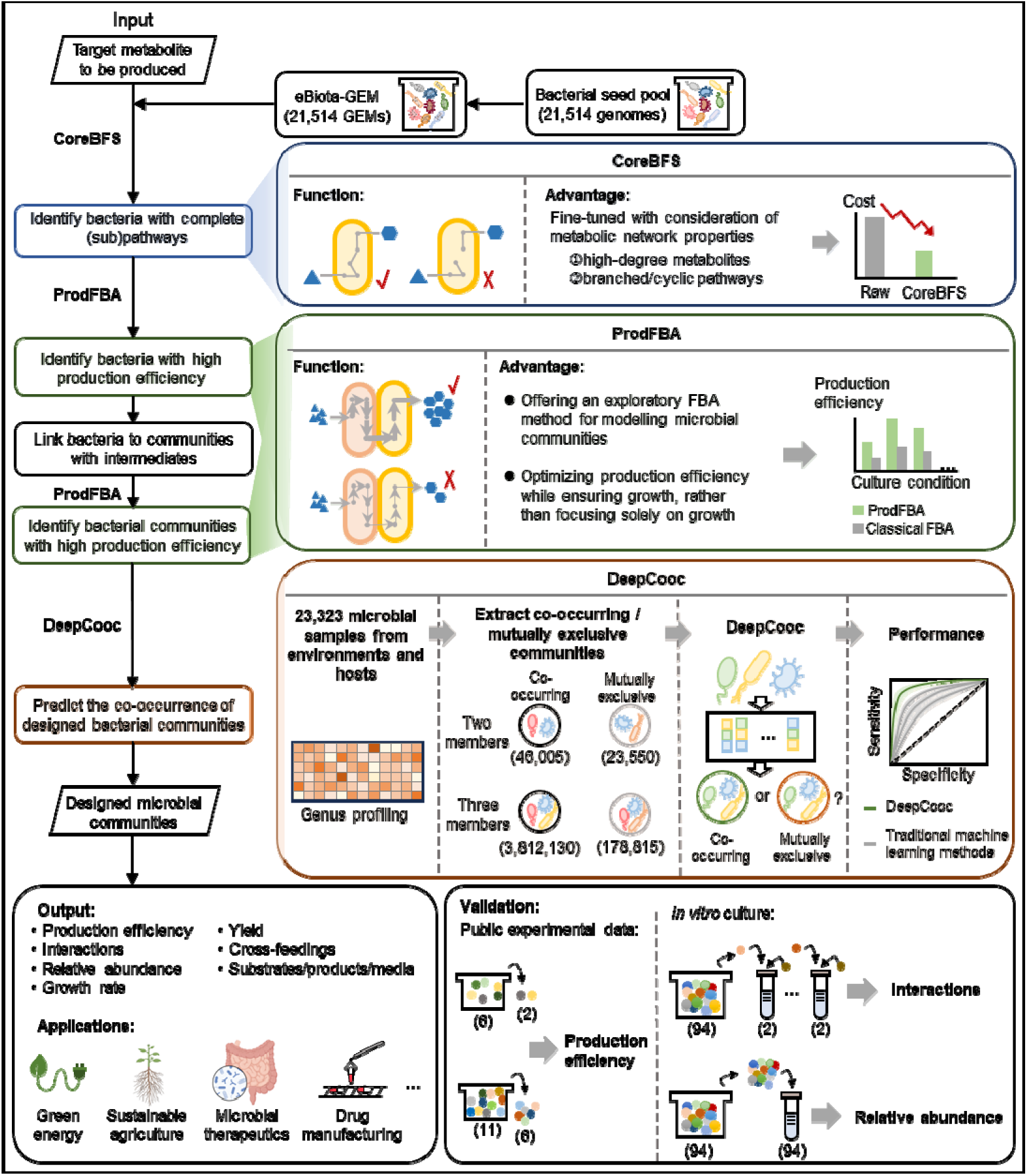
Workflow of eBiota. Overview of the eBiota pipeline and its three integrated algorithms: CoreBFS, ProdFBA, and DeepCooc. Numbers in parentheses represent the number of strains/cases. Production efficiency is shown as an illustrative example.

To build a large seed pool, we constructed eBiota-GEM, a GEM database comprising 21,514 bacterial genomes (see **Methods**, **Fig. 2A**, **Supplementary Table S1,** https://zenodo.org/records/13895108). eBiota-GEM was evaluated using MEMOTE^37^ and compared with two high-quality GEM repositories, namely AGORA^38^ and BiGG^39^. eBiota-GEM exhibited comparably good quality, underscoring its reliability as a basis for community design (**Extended Data Fig. 1** and **2**). These 21,514 GEMs collectively encompass 1,488 metabolites, of which 496 involved in exchange reactions, suggesting the potential to exchange 496 metabolites in extracellular space.

**Fig. 2.**
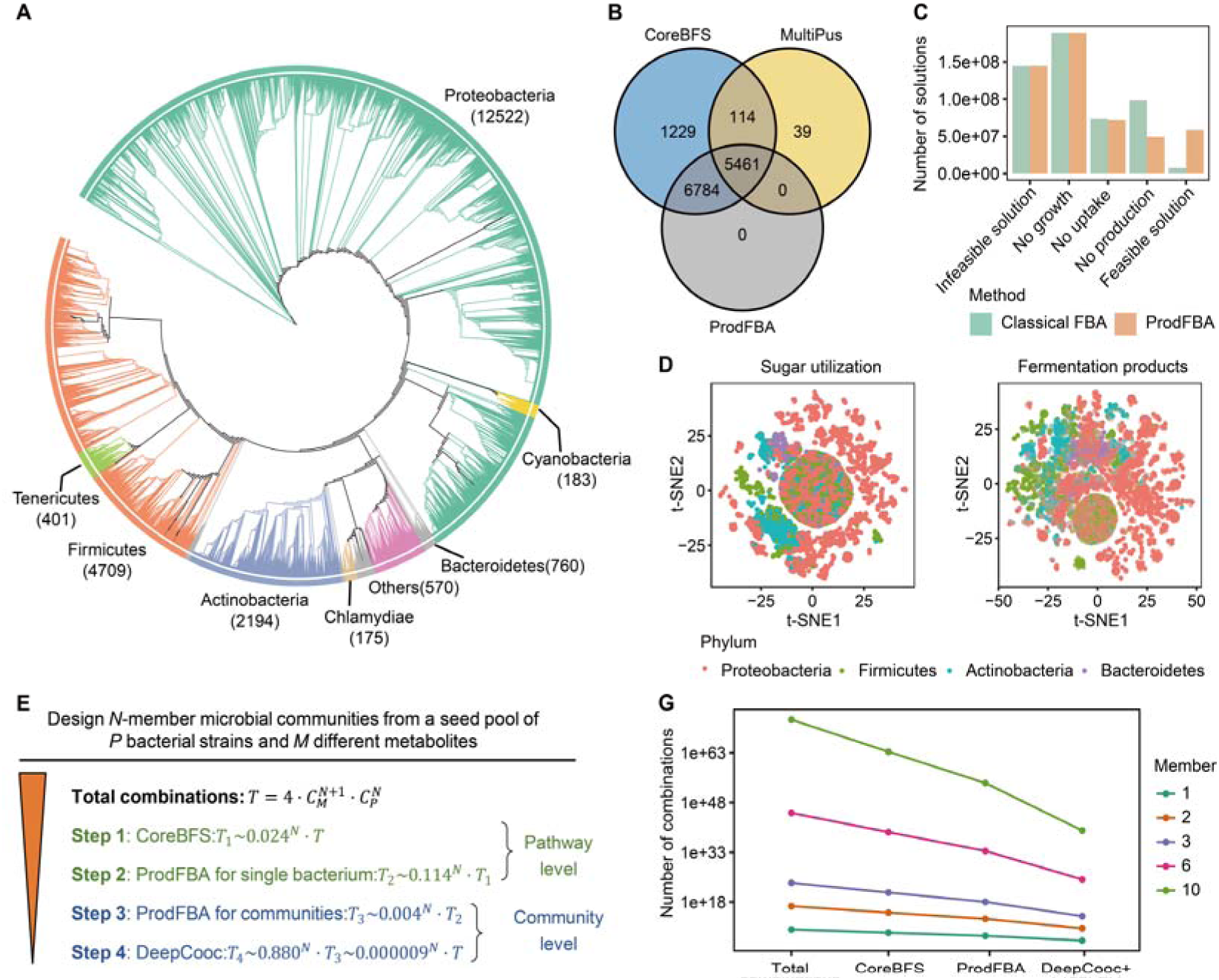
eBiota largely reduces the cost of designing microbial communities. (**A**) Phylogenetic tree constructed at the strain level based on the 16S rRNA gene sequences of each bacterium. Only 21,446 genomes with annotated 16S rRNA gene sequences are used. The numbers in parentheses indicate the number of genomes used in each phylum. (**B**) The Venn diagram shows the pathway calcualtes for hydrogen production using CoreBFS and MultiPus, with validation performed using ProdFBA. (**C**) Comparison of the solutions of classical FBA and ProdFBA. Infeasible: no flux distribution satisfies all constraints. No growth: a feasible solution exists, but the growth rate is zero. No uptake: a feasible solution exists and growth rate > 0, but uptake flux for the specified substrate is zero. No production: a feasible solution exists with growth rate > 0 and substrate uptake flux > 0, but secretion flux for the specified product is zero. Feasible: a feasible solution exists with growth rate > 0, substrate uptake flux > 0, and product secretion flux > 0 under the given conditions. (**D**) Visualizing the profiles of sugar utilization and fermentation products of microbial communities using t-SNE (t-distributed Stochastic Neighbor Embedding). Only the four phyla containing over 500 GEMs were shown. (**E**) Reductions in combinations achieved by eBiota. *T:* The total combinations*. T_1_, T_2_, T_3_, T_4_*: The number of remaining combinations after steps 1, 2, 3, and 4, respectively. (**F**) The number of combinations reduced by eBiota across microbial communities comprising 1, 2, 3, 6, and 10 members.

To rapidly determine whether a bacterium carries complete (sub)pathways that convert the substrate into the product, we developed CoreBFS, a graph-based breadth-first search (BFS) method tailored to metabolic networks. Unlike classical BFS, CoreBFS accounts for the scale-free and small-world properties of metabolic networks^40–42^, enabling effective handling of high-degree metabolites as well as branched and cyclic pathways (see **Methods**). We benchmarked CoreBFS against MultiPus^18^ using hydrogen production. Validation was performed with an FBA-based approach, in which the presence of a positive production flux necessarily indicates that the corresponding GEM contains complete biosynthetic pathways. Under conditions with all substrates supplied, CoreBFS achieved an accuracy of 0.938, identifying 13,588 candidate hydrogen producers, 12,245 of which were validated, with no false negatives (see **Methods**, **Fig. 2B**, **Supplementary Table S2**). In contrast, MultiPus achieved only an accuracy of 0.678, with 5,614 candidates identified, 153 false positives generated, and 6,784 true producers left undetected. These results established CoreBFS as a precise and high-throughput screening tool for the preliminary identification of functional bacterial candidates.

To quantitatively simulate individual strains and microbial communities under various conditions, rather than merely assessing pathway completeness, we developed ProdFBA (see **Methods**). ProdFBA combines the advantages of pFBA and FVA through multi-step optimizations: (i) calculating the maximum community growth rate using multi-compartment modeling, (ii) maximizing target production flux while maintaining ≥ 99% of maximal growth rate, and (iii) minimizing total flux. ProdFBA the simultaneous optimizes production or degradation efficiency and growth rate, overcoming the limitations of classical FBA, which focuses on growth rate maximization but fails to optimize production efficiency, particularly those that are uncoupled from growth^25,43^. In simulations of individual strains within eBiota-GEM, ProdFBA identified a higher proportion of bacteria with both high production efficiency and high growth rate (ProdFBA: 11.380%; Classical FBA: 1.515%) and a lower proportion of bacteria characterized by high growth rates but negligible production (ProdFBA: 9.625%; Classical FBA: 19.141%) (**Fig. 2C**). To validate the reliability of ProdFBA in capturing metabolic phenotypes, we benchmarked its predicted substrate uptake and product secretion profiles against curated data from 773 gut strains in the AGORA^38^. ProdFBA successfully reproduced 23 of 26 uptake reactions and 12 of 16 secretion reactions (**Extended Data Fig. 3C and D**), demonstrating its reliability in capturing microbial metabolic behaviors. Finally, taxonomic profiling of sugar utilization and fermentation products revealed significant variation across phyla (**Fig. 2D**, PERMANOVA, sugar utilization: R^2^ = 0.09, *p* < 0.001; fermentation products: R^2^ = 0.13, *p* < 0.001), a finding consistent with the functional specialization typically observed across bacterial phyla.

To infer co-occurrence patterns and gain insights into microbial coexistence, we implemented DeepCooc, a ResNet-based deep learning method that overcomes the degradation problem in deep neural networks^44–46^ (see **Methods**, **Extended Data Fig. 4**). Gold-standard labels were derived using HiOrCo^30^ from 23,323 samples in the Earth Microbiome Project (EMP)^47^, yielding 46,005 co-occurring and 23,550 mutually exclusive pairs, as well as 3,812,130 co-occurring and 178,815 mutually exclusive triplets. Using 335 transport reactions from GEMs as input features, DeepCooc achieved balanced accuracy scores of 0.966 for pairs and 0.911 for triplets, with AUROC (Area Under the Receiver Operating Characteristic Curve) values of 0.993 in both cases. These results outperformed traditional machine learning models, with improvements of at least 0.247 in balanced accuracy and 0.140 in AUROC (**Table 1**, **Fig. 3A** and **B**). To evaluate model robustness, we examined the effect of dropping features, reflecting incomplete or inaccurate metabolic annotations^48^. When randomly removing 1-300 of the 335 transport features, DeepCooc consistently maintained superior performance than other models (**Fig. 3C** and **D**, **Extended Data Fig. 5)**. Notably, AUROC values remained above 0.950 even with about 80 dropped features and showed small variation, indicating that model performance is not driven by any single dominant feature. These results demonstrate that DeepCooc is both highly accurate and robust in predicting microbial co-occurrence patterns.

**Fig. 3.**
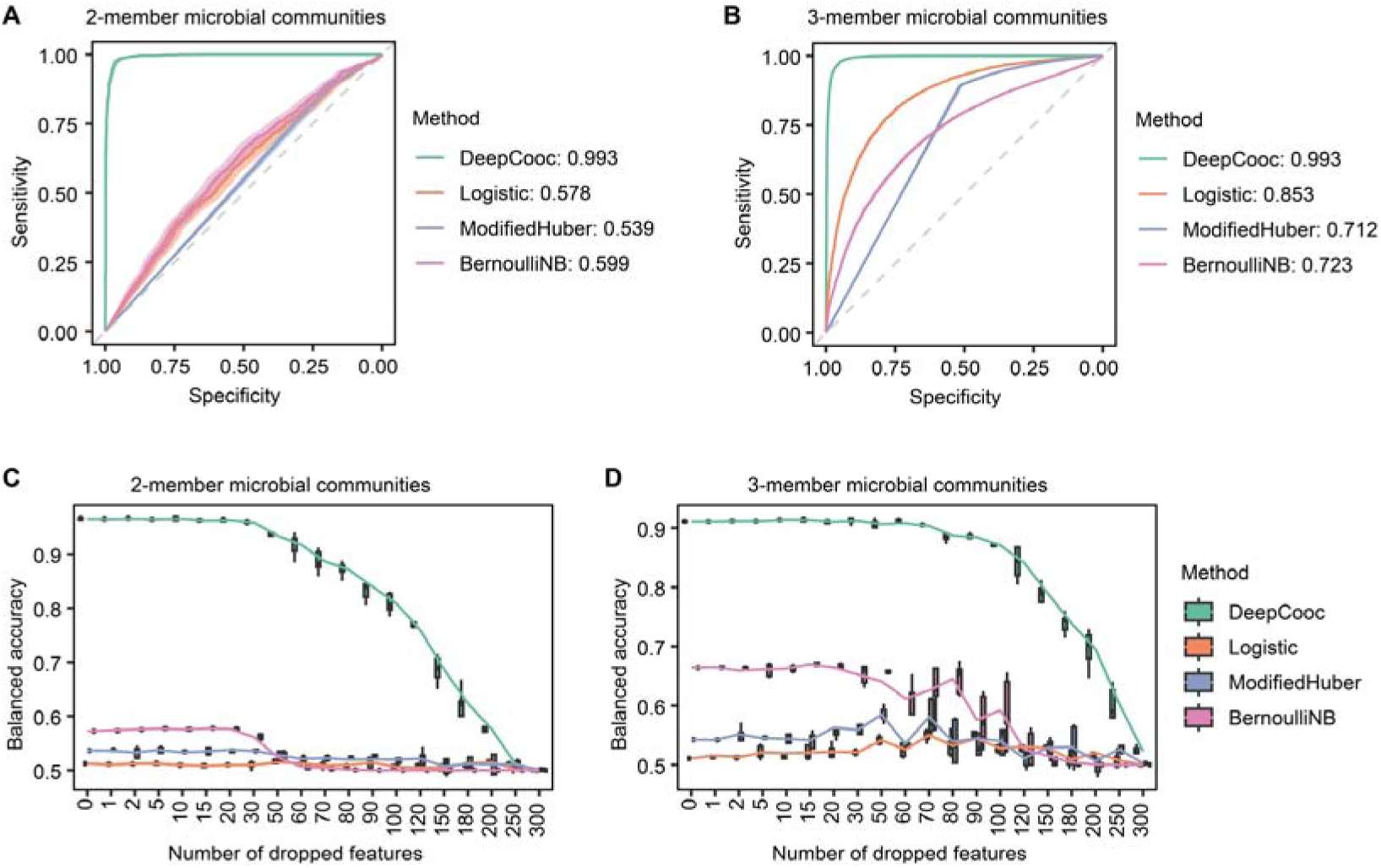
Comparison of performance of DeepCooc with other methods. Receiver operating characteristic curves for DeepCooc and other three methods applied to microbial communities consisting of two (**A**) and three (**B**) members. AUROC scores are presented in the legend. Changes in the balanced accuracy of four models when varying the number of features dropped, applied to microbial communities consisting of two (**C**) and three (**D**) members. The total number of features is 335. logistic: logistic regression; SVM: support vector machine. Naive Bayes: Bernoulli naive bayes.

**Table 1.** Comparison of the performance of DeepCooc and other models on the test dataset.

| Members in community | Model | F1 score | Balanced accuracy | AUROC |
| --- | --- | --- | --- | --- |
| 2 | DeepCooc | **0.978** | **0.966** | **0.993** |
|  | Logistic | 0.791 | 0.512 | 0.578 |
|  | SVM | 0.774 | 0.537 | 0.539 |
|  | Naive Bayes | 0.709 | 0.572 | 0.599 |
| 3 | DeepCooc | **0.995** | **0.911** | **0.993** |
|  | Logistic | 0.977 | 0.511 | 0.853 |
|  | SVM | 0.975 | 0.542 | 0.712 |
|  | Naive Bayes | 0.843 | 0.664 | 0.723 |

*Note:* Logistic: logistic regression; SVM: support vector machine. Naive Bayes: Bernoulli naive bayes.

Then we used eBiota to design one- and two-member communities from eBiota-GEM, maximizing production efficiency across all potential target metabolites (**Supplementary Table S3**, https://zenodo.org/records/13895108). For the *de novo* design of *N*-member microbial communities selected from a pool of 21,514 taxa and 496 potential substrates/products under four conditions, there are 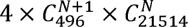 possible combinations. For N = 2, this corresponds to approximately 1.871× 10^16^. eBiota reduces the combinations to 0.000009*^N^* of the total. Specifically, sequential filters by CoreBFS, ProdFBA at the single-microbe level, ProdFBA at the community level, and DeepCooc progressively reduced the design space to 0.024*^N^*, 0.114*^N^*, 0.004*^N^*, and 0.880*^N^*of the previous stage, respectively **(Fig. 2E** and **F**). The resulting 18,808,505 communities generate 247 products, mainly amino acids, followed by organic acids, sugars, and lipopolysaccharides. They are also capable to utilize 309 substrates, most commonly sugars and organic acids, followed by polysaccharides and amino acids. A detailed comparison between eBiota and existing tools is provided in **Table 2**. In summary, eBiota is functionally comprehensive, computationally efficient, and uniquely enables the *ab initio* design of microbial communities from large-scale seed pools.

**Table 2.**
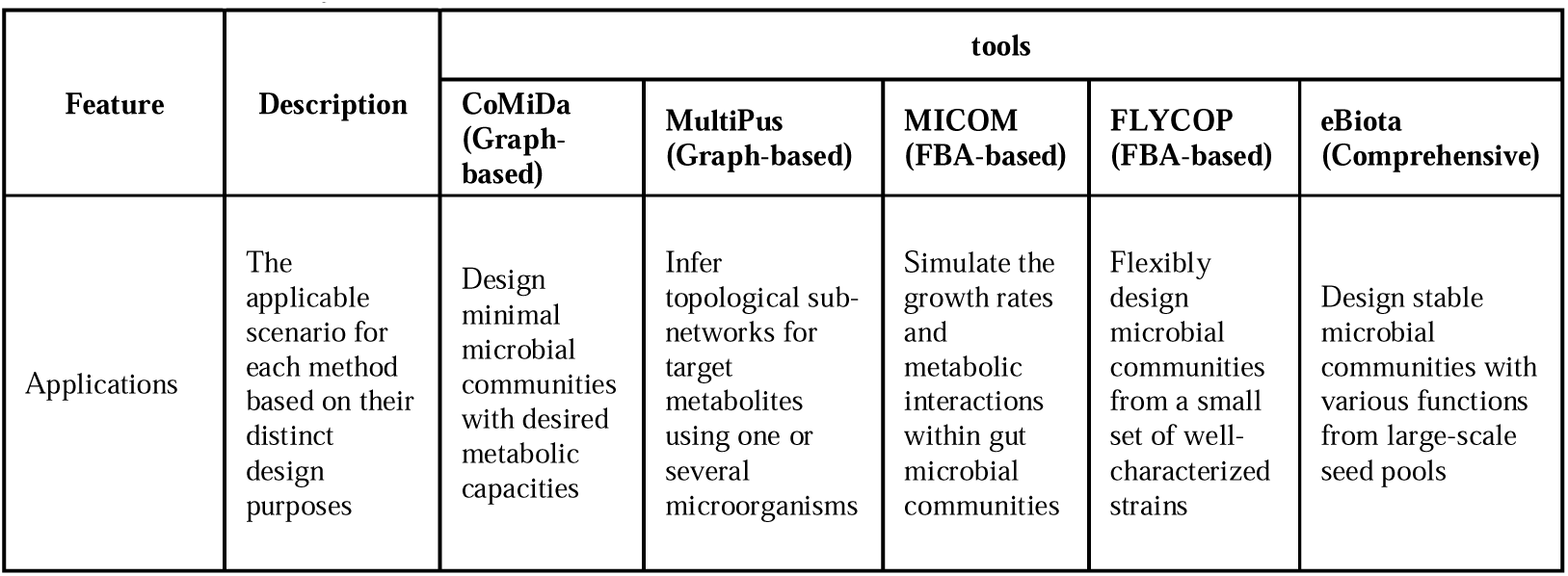
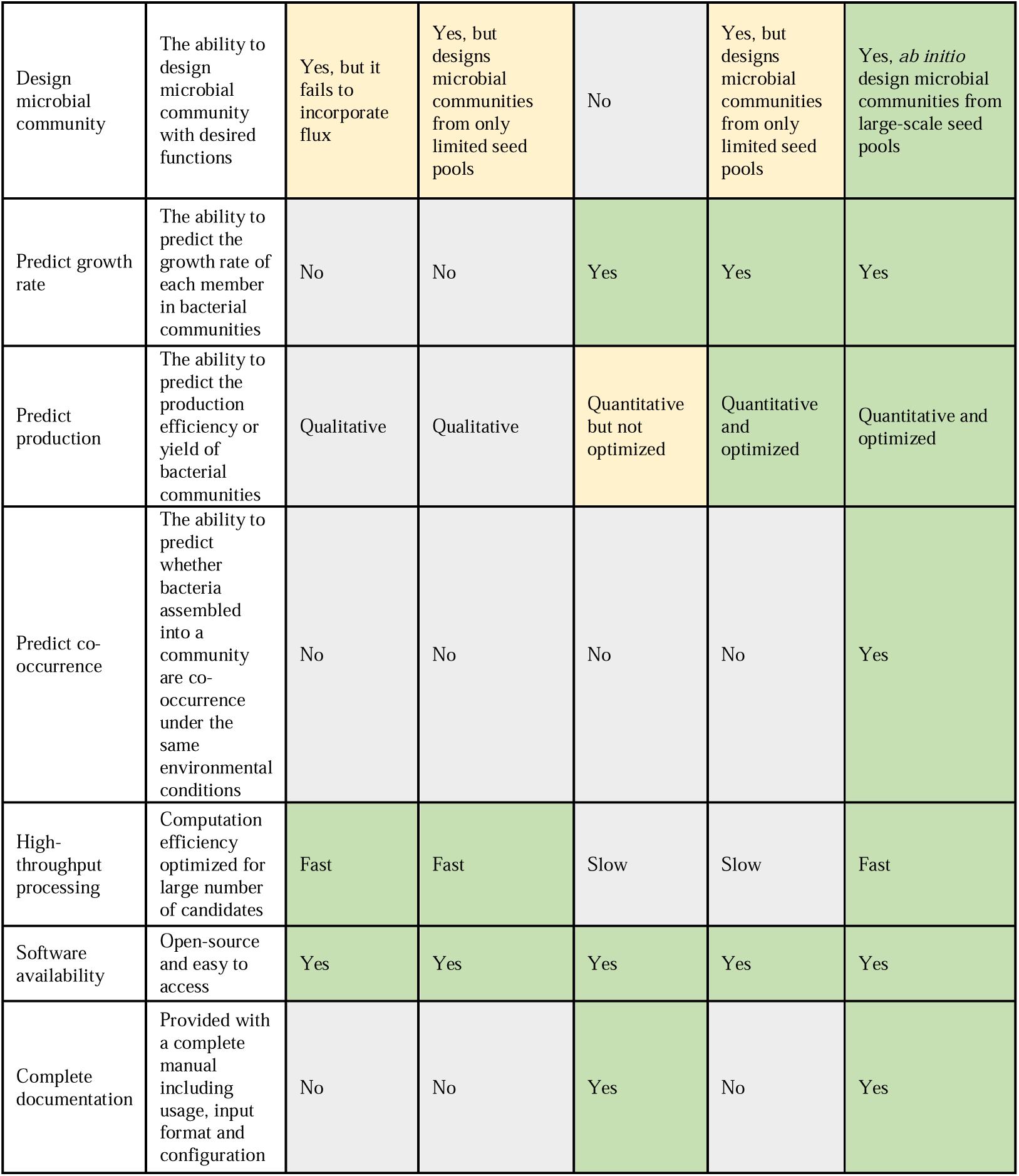
Summary of the features of eBiota and relevant tools.

### eBiota accurately designs microbial communities having the capability to produce different metabolites

Extensive experimental data on microbial production of hydrogen, short-chain fatty acids (SCFAs), and amino acids provide a valuable opportunity to validate eBiota. We employed eBiota to design microbial communities optimized for hydrogen production, while also extending the analysis to SCFAs and amino acids across systems ranging from mono-cultures to multi-member consortia. The simulation results were compared with publicly available experimental data to evaluate eBiota’s *ab initio* design capability and its accuracy in reproducing community metabolic behaviors.

To examine eBiota’s performance in calculating production within single-taxon systems, we compared its outputs with experimental data on hydrogen and two SCFAs (formate and acetate) from 24 genera summarized by Djemai *et al.* across more than 100 studies^49^. eBiota’s simulations showed strong agreement with experimental observations, achieving F1-score of 0.839, 0.757, and 0.909 for hydrogen, formate, and acetate, respectively (**Fig. 4A**), outperforming MICOM—which yielded F1-scores of 0.727, 0.743, and 0.857 and is itself a widely used microbial community modeling tool that ranks among the top methods in quantitative assessment^50^. Moreover, genera reported as hydrogen producers exhibited significantly higher efficiency rankings in eBiota (150.154 ± 108.163) than those without such reports (574.375 ± 168.280, Wilcoxon rank-sum test, two-tailed *p* = 0.0004; **Fig. 4B**). Notably, eBiota identified the anaerobic moderately thermophilic bacterium *Tenuifilum thalassicum* 38H-str as the most efficient hydrogen producer among 21,514 taxa when utilizing polymeric substrates such as starch and amylose (**Supplementary Table S4**). Its efficiency declined sharply on simple sugars like glucose, consistent with previous studies^51,52^ showing that thermophilic bacteria achieve near-theoretical hydrogen yields on polymeric substrates but exhibit poor growth on simple sugars. These results demonstrate that eBiota effectively captures production capabilities of individual strains.

**Fig. 4.**
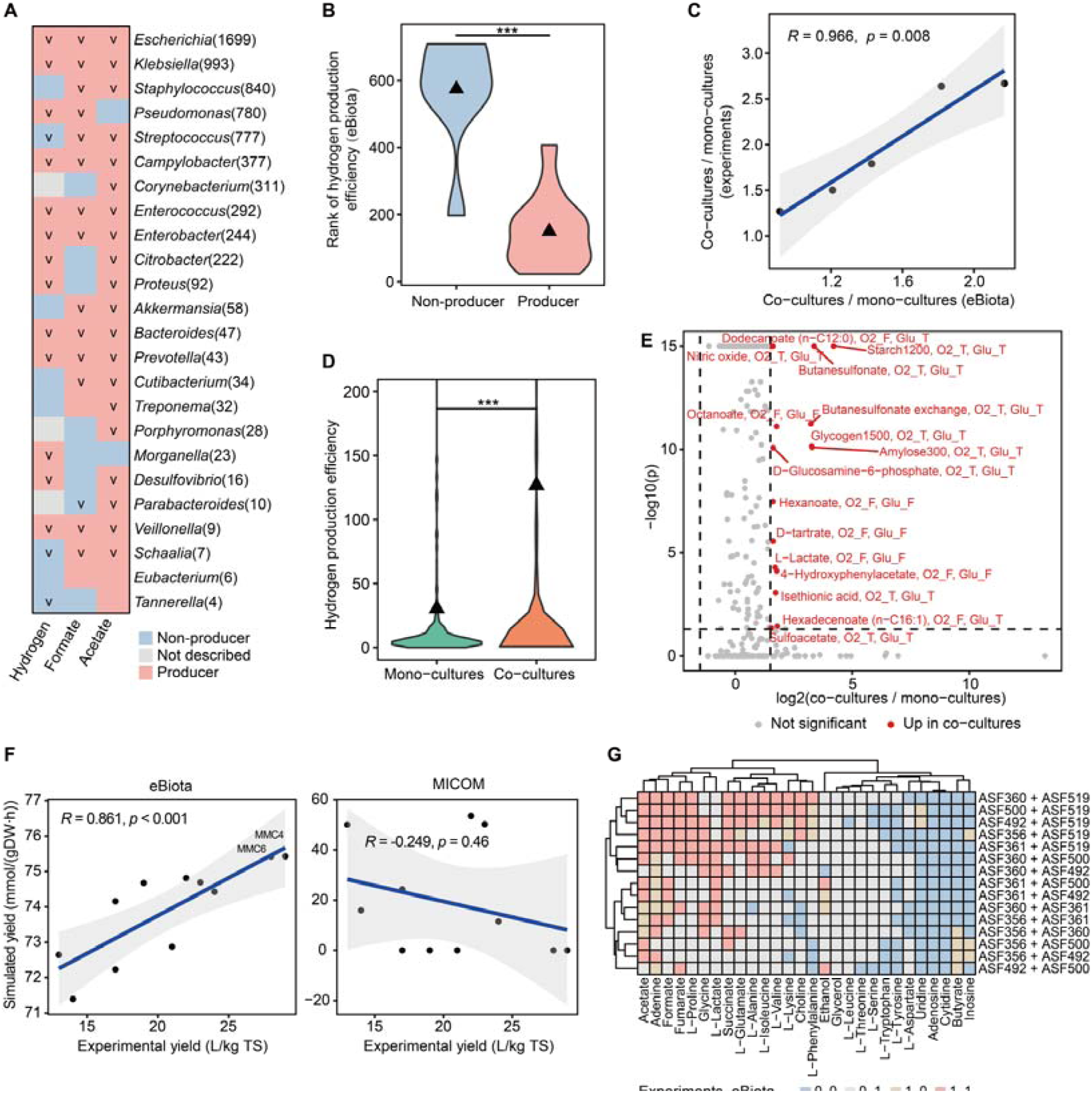
Comparison of eBiota-designed microbial communities and experimental data. (**A**) Heatmap indicates whether each production capability is documented in the review. A “v” in the heatmap denotes capabilities that were accurately predicted by eBiota. The numbers in parentheses represents the number of GEMs in each genus. (**B**) Comparison of ranked hydrogen production efficiency calculated by eBiota for genera documented as hydrogen producers (*n* = 13) versus those that are not (*n* = 8). Lower rank indicates higher hydrogen production efficiency. Triangles indicate the mean value for each group. ***: *p* < 0.001, Wilcoxon rank-sum test. (**C**) Relationship between hydrogen production efficiency of co-cultures and mono-cultures. The horizontal axis represents the relationship calculated by eBiota, while the vertical axis shows the experimental data. Pearson correlation coefficient, *p* value and a linear regression line with 95% confidence intervals are shown. Comparisons with MICOM were not feasible because, except for case 33, it predicted zero product fluxes in either mono- or co-cultures. (**D**) Comparison of hydrogen production efficiency between mono-cultures and co-cultures calculated by eBiota (*n* = 308 for both). ***: *p* < 0.001, Wilcoxon signed-rank test. (**E**) Volcano plot showed the differences in hydrogen production efficiency between mono-cultures and co-cultures calculated by eBiota. Differences are considered statistically significant when the *p* < 0.05 and |log2(hydrogen production efficiency of co-cultures / hydrogen production efficiency of mono-cultures)| >1.5. O2_T, Glu_T: With both oxygen and glucose; O2_F, Glu_T: With oxygen only; O2_T, Glu_F: With glucose only; O2_F, Glu_F: With neither oxygen nor glucose. (**F**) Comparison of the maximum hydrogen yield calculated by eBiota and MICOM against publicly available experimental results. Hydrogen yield refers to the production of hydrogen generated per unit of substrate. Spearman correlation coefficient, *p* value and a linear regression line with 95% confidence intervals are shown. TS: Total solids of pea-shell, the substrate in the experiment. DW: Dry weight. **(G)** Comparison of production capability of 28 metabolites generated by microbial communities designed by eBiota against publicly available experimental results. Blue: both experimental and eBiota results are 0 (no production capability). Red: both are 1 (production capability observed experimentally and predicted by eBiota). Gray: experimental result is 0 but eBiota result is 1 (predicted only). Yellow: experimental results are 1 and eBiota results are 0 (observed experimentally only). ASF356: *Clostridium* sp.; ASF360: *Lactobacillus intestinalis*; ASF361: *Lactobacillus murinus*; ASF492: *Eubacterium plexicaudatum*; ASF500: *Pseudoflavonifractor* sp.; ASF519: *Parabacteroides goldsteinii*.

To evaluate eBiota’s capability for community design and hydrogen production in multi-member systems, we conducted an extensive literature review and identified 34 two-member and nine three-member hydrogen-producing consortia. Among these, eBiota successfully simulated hydrogen production and co-occurrence in 29 of the 34 two-member communities (or 17 of 17 when excluding species with no more than three GEMs) and in all nine three-member communities (**Supplementary Table S5** and **S6**). In contrast, MICOM successfully simulated 23 of the 34 two-member communities but only 3 of the nine three-member communities. Moreover, eBiota reproduced three representative hydrogen production processes^53,54^. The first is dark fermentation, where acidogenic bacteria such as *Clostridium*, *Enterobacter*, and *Bacillus*, along with several thermophilic species, ferment organic substrates to generate hydrogen with volatile fatty acids and alcohols as by-products. In eBiota, a co-culture of *Citrobacter freundii* and *Clostridium butyricum* produced hydrogen from organic substrates with acetate, CO□, ethanol, and lactate as by-products. eBiota also simulated a mixed culture of *Acetivibrio thermocellus*, *Thermoanaerobacterium aotearoense*, and *Saccharomyces cerevisiae*, identifying glucose and xylose as optimal substrates for maximal hydrogen yields, consistent with experimental results^55^. The second process, dark-photo co-fermentation, involves cooperation between photosynthetic bacteria (*Rhodopseudomonas*, *Rhodobacter*) and dark fermentative bacteria to further convert volatile fatty acids and other intermediates generated during dark fermentation. eBiota reproduced this process in a co-culture of *Clostridium beijerinckii* and *Rhodobacter sphaeroides*, which utilized complex substrates (amylose, glycogen, starch) to produce hydrogen through cross-feeding of glucose and by-products including acetate, CO□, ethanol, and lactate. eBiota also simulated hydrogen production by *Klebsiella pneumoniae* and *Rhodospirillum rubrum*, consistent with observations that *R. rubrum* contaminated with *K. pneumoniae* can degrade cellulose to generate hydrogen^56^. The third process, interspecies electron transfer, exemplified by *Geobacter metallireducens* and *Geobacter sulfurreducens*, was also reproduced by eBiota through cross-feeding of H□ and Fe²□, aligning with experimental observations^57^.

To further examine eBiota’s accuracy in comparing hydrogen production efficiencies between mono- and co-cultures, we focused on five of the 17 two-member communities for which experimental data confirmed hydrogen production by both constituent strains in mono-culture. eBiota accurately reproduced the relative efficiencies of individual strains and their corresponding co-cultures, showing strong correlation with experimental data (**Fig. 4C**, Pearson correlation: *R* = 0.966, *p* = 0.008) whereas MICOM successfully predicted only a single case. We then extended this comparison to all one- and two-member communities designed by eBiota using the eBiota-GEM dataset. Under identical culture conditions and substrates, co-cultures exhibited significantly higher hydrogen production efficiencies than mono-cultures, with a median 1.5-fold increase (IQR: 1.149-2.069; **Fig. 4D**, Wilcoxon signed-rank test, two-tailed *p* < 0.001), and no mono-culture outperformed its corresponding co-culture (**Fig. 4E**). Co-cultures with greater efficiency gains tended to utilize substrates with higher carbon numbers, such as polysaccharides and medium-chain fatty acids, and showed positive correlation with metabolic cross-feeding (**Extended Data Fig. 6**, Pearson correlation: *R* = 0.683, *p* = 0.004 for carbon count; *R* = 0.554, *p* = 0.026 for cross-feeding). These findings are supported by previous experimental evidence showing that co-cultures degrade complex substrates and exchange metabolites more effectively than mono-cultures^53,58^.

To assess eBiota’s performance in designing larger microbial communities and estimating their hydrogen yields, we used the experimental dataset from Patel *et al.*^59^, which measured hydrogen yields for 11 mixed microbial cultures (MMC1–MMC11), each comprising six strains. MMC4 and MMC6 exhibited the highest yields, with MMC4 slightly outperforming MMC6. Based on this dataset, we constructed a bacterial seed pool of 11 taxa and used eBiota to design microbial communities from this pool (**Supplementary Table S7**). eBiota successfully identified hydrogen production capabilities across all 11 mixed cultures. The estimated hydrogen yields correctly ranked MMC4 highest, followed by MMC6, consistent with experimental observations. Furthermore, hydrogen yields calculated by eBiota correlated strongly with experimental data (**Fig. 4G**, Spearman correlation: *R* = 0.861, *p <* 0.001). In contrast, the yields computed using MICOM showed no significant correlation with the experimental data. These results demonstrate that eBiota effectively designs multi-taxa microbial communities from a given seed pool and accurately reproduces their hydrogen yield.

Finally, to evaluate eBiota’s capacity to design communities for the production of multiple metabolites, we employed a published experimental dataset that measured the output of 28 metabolites from pairwise co-cultures of six altered Schaedler flora (ASF) strains isolated from the mouse gut^60,61^. These metabolites included amino acids, SCFAs, other organic acids, nucleosides, and vitamins. Using these six strains as a seed pool, eBiota was applied to design all possible two-member communities and simulate their metabolite production capabilities. The results closely matched experimental data, with all designed communities exhibiting co-occurrence and achieving a recall of 0.770 across the 28 metabolites (**Fig. 4F**). Some metabolite productions estimated by eBiota but not reported experimentally likely reflect the its ability to identify communities with untested production potential. In contrast, MICOM failed to reproduce many experimentally observed production capabilities, each achieving a recall of only 0.361 (**Supplementary Table S8**). These results demonstrate that eBiota reliably designs microbial communities and simulates the production of a wide range of metabolites.

### eBiota simulated community interactions and composition *in vitro*

To assess eBiota’s ability to design microbial communities and simulate microbial interactions, we conducted *in vitro* culture experiments. Plant pathogens cause major agricultural and economic losses and their control is often hindered by poor understanding of interactions with root-associated microbiomes, which are increasingly recognized as drivers of natural pathogen resistance^62,63^. To explore this, we used 94 bacterial strains isolated from plant roots together with the pathogen *R. solanacearum* QL-Rs1115^64^ (see **Methods**, **Supplementary Table S9**). The growth of *R. solanacearum* QL-Rs1115 was measured in both mono-culture and sequential culture with each strain to quantify experimental competition intensity, while eBiota was used to simulate these communities and calculate pathogen growth *in silico*. Strains were classified as competitors if simulated growth of *R. solanacearum* QL-Rs1115 in sequential culture decreased by at least 10% relative to mono-culture, a commonly applied threshold for detecting microbial interaction^65,66^. eBiota identified 88 of the 94 strains as competitors, which exhibited significantly higher experimentally measured competition intensity than non-competitors (**Fig. 5A**; Wilcoxon rank-sum test, two-tailed *p* = 0.015). As a case study, we further examined strains showing the greatest and the least effects on the growth of *R. solanacearum* QL-Rs1115 in sequential culture experiments and used eBiota to perform Pareto optimality to delineate the non-dominated trade-off frontier among their growth. Among the five strains with the greatest effects, four were identified by eBiota as engaging in strong competition with the pathogen, as reflected by Pareto fronts showing minimal mutual benefit and marked inhibition of pathogen growth (**Fig. 5B**). In contrast, among the five strains with the least effects, three exhibited nearly independent growth from *R. solanacearum* QL-Rs1115 on the Pareto front. These results confirm eBiota’s ability to model microbial interactions.

**Fig. 5.**
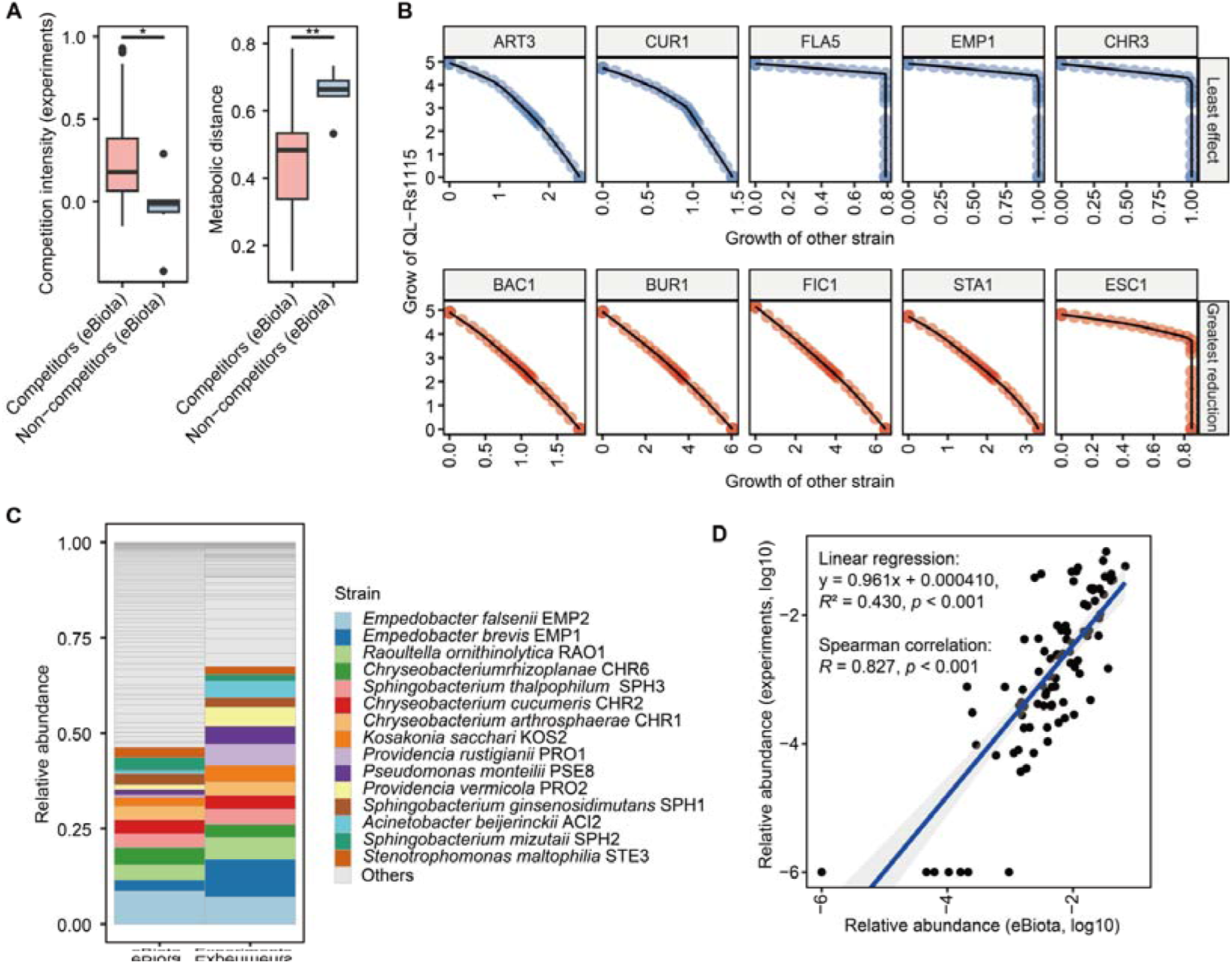
Comparison of eBiota-designed microbial communities with the *in vitro* culture experiments. (**A**) Left: The competition of each of 94 strains against *R. solanacearum* QL-Rs1115 in *in vitro* culture experiments was compared with the results calculated by eBiota (competitors: *n* = 88; non-competitors: *n* = 6). Right: Comparison of the metabolic distance from *R. solanacearum* QL-Rs1115 for competitors and non-competitors. *: *p* < 0.05, **: *p* < 0.01, Wilcoxon rank-sum test. (**B**) The Pareto front illustrates the trade-off between the growth of *R. solanacearum* QL-Rs1115 and that of the interacting strain. The top five strains with the highest and lowest effect on *R. solanacearum* QL-Rs1115 are shown. (**C**) Comparison of the mean relative abundances of 94 strains calculated by eBiota and those observed in *in vitro* co-culture experiments. (**D**) Relationship between the mean relative abundances of 94 strains calculated by eBiota and those observed in *in vitro* co-culture experiments. Linear regression equations, *R*^2^, and *p* value, along with the Spearman correlation coefficient and *p* value are shown. QL-Rs1115: *R. solanacearum* QL-Rs1115; CUR1: *Curtobacterium citreum*; FLA5: *Flavobacterium naphthae*; EMP1: *Empedobacter brevis*; ART3: *Arthrobacter globiformis*; CHR3: *Chryseobacterium daecheongense*; BAC1: *Bacillus amyloliquefaciens*; BUR1: *Burkholderia cenocepacia*; FIC1: *Fictibacillus barbaricus*; STA1: *Staphylococcus hominis*; ESC1: *Atlantibacter* (formerly *Escherichia*) *hermannii.*

To investigate the mechanisms underlying these interactions, we applied a phylogenetically adjusted quantification method^67,68^ to calculate metabolic competition and complementarity indices between each strain and *R. solanacearum* QL-Rs1115. Metabolic competition represented the overlap in metabolic reactions between GEMs, whereas complementarity reflected the potential to utilize each other’s by-products. We also calculated the composite metabolic distance score, defined as 1-□(competition index□-□complementarity index)^67^, to represent the overall functional dissimilarity. eBiota was further used to estimate potential cross-feedings and co-occurrence patterns. The results showed that competitors had significantly lower metabolic distance scores and higher competition indices compared to non-competitors (**Fig. 5A** and **Extended Data Fig. 7**; Wilcoxon rank-sum test, both two-tailed *p* < 0.05), while no significant differences were observed in complementarity indices or the number of cross-feedings. All non-competitors were predicted to co-occur with *R. solanacearum* QL-Rs1115, whereas seven competitors were predicted to be non-co-occurring and showed even lower metabolic distance scores and higher competition indices (**Supplementary Table S9**; Wilcoxon rank-sum test, two-tailed *p* < 0.05). Altogether, the observation that most strains competed with *R. solanacearum QL-Rs*1115 suggests that habitat filtering, which promotes the coexistence of functionally similar taxa, may be a major driver of colonization and assembly in the root microbiome, while the partial non-co-occurrence of competitors with the highest functional similarity indicates that excessive overlap could also lead to exclusion.

To test eBiota’s ability in calculating bacterial composition within complex multi-species communities, we combined all 94 strains into a single community and quantified their relative abundances using 16S rRNA gene sequencing (see **Methods**). eBiota successfully simulated the community structure of this 94-member consortium, showing a strong correlation with experimentally measured mean relative abundances (**Fig. 5C** and **D**, linear regression: y = 0.961x + 0.000410, 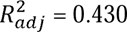, *p* < 0.001; Spearman correlation: *R* = 0.827, *p* < 0.001) and a low Bray-Curtis distance of 0.386. These results demonstrate that eBiota can effectively simulate and design microbial communities comprising large numbers of taxa.

## Discussion

The rational design of microbial communities with defined metabolic functions is a long-standing goal in synthetic ecology and microbiome engineering. Existing approaches often rely on descriptive modeling or predefined small seed pools, which severely limit their capacity to explore the vast microbial diversity. Expanding the design space to large microbial pools can uncover new functional combinations and open opportunities for discovering highly efficient consortia. Here, we present eBiota, a platform that integrates rapid optimization algorithms and deep learning to enable *ab initio* design of microbial communities with desired functions from large seed pools.

eBiota aims to address two key challenges in the rational design of microbial communities: the combinatorial explosion of possible community configurations and the difficulty of identifying both functionally efficient and ecologically compatible members. To overcome these challenges, eBiota adopts a hierarchical screening strategy based on three purpose-built modules. The graph-based CoreBFS module considers only the topological structure of metabolic networks and thus rapidly identifies taxa that together complete the desired metabolic pathways. The ProdFBA module applies flux-based optimization that integrates metabolic and environmental constraints to identify strains and communities capable of maintaining growth rate while maximizing production efficiency. The third module, DeepCooc, is a deep learning model trained on over 23,000 real-world microbiome samples that infers pairwise and higher-order microbial co-occurrence patterns. As summarized in **Table 2**, eBiota is, to the best of our knowledge, the only approach that integrates pathway-based screening, flux optimization, and data-driven ecological inference within a unified platform, and the only one capable of designing productive and ecologically relevant communities from large microbial pools.

As a one-stop *in silico* microbial laboratory, eBiota constructs digital twins of microbiome and is able to predict diverse microbial behaviors, including production or degradation efficiency, metabolic interactions, cross-feeding relationships, and community composition. Detailed protocols and use cases are available on our website (https://github.com/allons42/eBiota). In addition, eBiota can reveal untested metabolic potential by identifying possible products and substrates beyond those previously confirmed experimentally, such as in the examples of eBiota-GEM and ASF strain simulations, although some maybe false positives. Furthermore, it can compare community performance under different culture conditions, as shown by its accurate prediction of the relative efficiencies of individual strains and their co-cultures, and can provide mechanistic insights, as shown by reproducing known hydrogen production processes and modeling microbial competition with pathogens. As an early exploration of artificial intelligence-aided microbial communities design, eBiota is expected to serve as a versatile platform for future applications in green energy production, sustainable agriculture, microbiota-based therapies, pharmaceutical synthesis, industrial fermentation, and environmental remediation.

Despite its broad applicability, eBiota has several limitations. First, it currently uses automatically reconstructed GEMs; incorporating curated models such as AGORA2 and integrating experimental data will improve its accuracy and reliability. Second, the current version includes only bacterial genomes; extending it to fungi and archaea will yield a more comprehensive representation of natural microbial ecosystems. Third, like most static FBA approaches, ProdFBA simulates steady-state behavior. Incorporating dynamic flux balance analysis or reinforcement learning could enable eBiota to capture adaptive and evolutionary processes in microbial communities. Fourth, DeepCooc predicts co-occurrence from real-world microbiome data but does not directly model stable coexistence. The accumulation of larger experimental datasets and explicit modeling of environmental, spatial, and temporal factors will further enhance eBiota’s ability to predict stable coexistence. Finally, eBiota follows a bottom-up design strategy; integrating top-down approaches could better link designed and natural communities, enhancing both predictive realism and experimental relevance^69^.

## Methods

### GEM reconstruction and phylogenetic tree analysis

We downloaded protein sequences for 21,514 complete non-redundant bacterial genomes from RefSeq (https://www.ncbi.nlm.nih.gov/refseq/). We employed CarveMe (version 1.4.1)^70^ to reconstruct GEMs for each bacterial genome with the universal bacterial template. This template includes three compartments (cytosol, periplasm, and extracellular spaces), 5532 reactions (including 1870 transport and 645 exchange reactions). Transport reactions characterize the transport of metabolite across compartments and exchange reactions represent the flux of substrate uptake and product discharge. CarveMe is a fast-automated method for reconstructing GEMs that has been validated to perform comparably to the (semi-)manual methods in reproducing experimental phenotypes^70^. The quality of these GEMs was assessed using the MEMOTE^37^ test suite, a standardized method for quality control of GEMs.

We extracted the 16S rRNA gene sequences from 21,446 bacteria based on their annotations. Multiple sequence alignments of 16S rRNA gene sequences were constructed using MUSCLE (version 5.1.linux64)^71^ and the approximately maximum likelihood phylogenetic tree was built using FastTree (version 2.1.11)^72^ with the generalized time-reversible model.

### The annotation of bacteria

Standardized bacterial information, including isolation source and oxygen tolerance, was retrieved from BacDive^73^ (https://bacdive.dsmz.de/). Of the 21,514 strains in eBiota-GEM, 2,282 were matched to BacDive using NCBI assembly accession numbers, and 17,173 were matched through taxon IDs at the species level. Bacteria labeled “#Feces (Stool)” were classified as gut bacteria, totaling 7,499 strains across 280 species. Of these species, 127 were present in AGORA, corresponding to 6,142 strains. Oxygen tolerance annotations were available for 16,032 strains. Detailed information can be found in **Supplementary Table S1**.

### Implementation of CoreBFS

CoreBFS is a graph-based, breadth-first search algorithm for quickly identifying whether a bacterium has complete metabolic pathways from substrates (or intermediates) to products (or intermediates). Taking the search from intermediates to products as an example, we compile a list of potential products from the exchange reactions in the GEM and use this list as the starting point. When the searched metabolite is identified as a component of the medium or as a potential intermediate, the search is terminated, and the complete pathway from the intermediate to the product is recorded. CoreBFS was implemented in Perl (version 5.26.1).

Considering that the metabolic network is scale-free and exhibits small-world properties^40–42^, we have implemented the following strategies: 1) The degree distribution of a scale-free network follows a power law, meaning that most nodes have few connections, while a small number of nodes possess a disproportionately large number of connections. The latter, known as “currency metabolites” in metabolic networks, participate in numerous reactions and overshadow the structure of the GEM. They are often removed in other studies^74^ and are pre-stored in our CoreBFS. Specifically, metabolites with a degree greater than 20, such as metal ions, oxygen, adenosine triphosphate, and coenzyme A, are considered as “currency metabolites” in this study, as they are involved in large number of reactions but do not define or specify any individual pathways^74^. The pre-storage of “currency metabolites” effectively narrows the search scope. 2) The metabolic network is a small-world graph^40,75^, meaning it is highly clustered with a small average path length. Thus, a maximum reaction step limit of 30 was set for pathways, a threshold that has been shown to cover about 99% of pathways^76^ (**Extended Data Fig. 3A**). 3) Some pathways, such as glycolysis, the tricarboxylic acid cycle, the pentose phosphate pathway, and the Calvin cycle, which are common in many microorganisms, are pre-stored. 4) When a previously recorded metabolite is encountered again, the current search branch is pruned.

We compared the performance of CoreBFS with MultiPus^18^. MultiPus was implemented in Python 2.7 environment using Clingo4.5.4^77^ as the Answer Set Programming solver. CoreBFS and MultiPus were applied to identify hydrogen-producing bacteria across 21,514 bacteria in eBiota-GEM, with all substrates available. CoMiDA^17^, another graph-based method, was excluded due to the complexity of code implementation.

### The implementation of ProdFBA

We developed ProdFBA to quantitatively simulate the mono-culture of individual strains as well as co-cultures of microbial communities with high efficiency. ProdFBA is a multi-step linear programming algorithm that involves 1) maximizing production (or degradation) efficiency while maintaining at least 99% of the maximum growth rate, rather than maximizing growth rate as in classical FBA. Production (or degradation) efficiency is calculated as the sum of the maximum production (or degradation) flux multiplied by the growth rate of each bacterium, as described in a previous study^66^; and 2) minimizing total flux summation to reduce redundant pathways, similar to parsimonious enzyme usage FBA^23^. Therefore, ProdFBA can be formulated as follows:

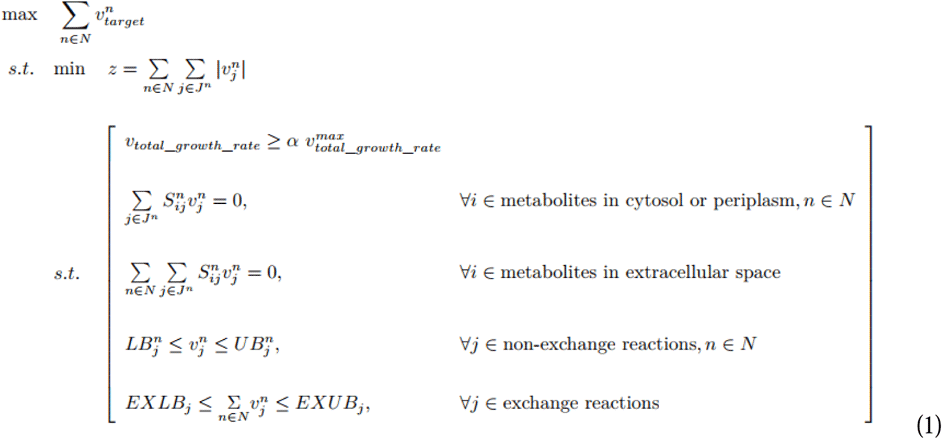

Where N is the GEM set of microorganisms in the community. J^n^ denotes the set of reactions in the GEM n. 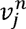 is the flux of reaction j in the GEM n. EXLB_j_ and EXUB_j_ are the lower bound and upper bound for exchange reaction j in the extracellular space, respectively. 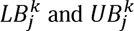 are the lower bound and upper bound for non-exchange reaction j in the GEM n, respectively. S^n^ is the stoichiometry matrix of GEM n. The parameter a is set to 0.99 to ensure that the growth rate does not fall below 99% of the maximum growth rate.

Users have the flexibility to define their preferred growth medium. We utilized Luria-Bertani (LB) medium for the initial design, which is one of the most basic media frequently used in simulated cultures studies. We supplement the medium with basic nutrients such as amino acids, vitamins, and nucleotides (**Supplementary Table S3**). To support the growth of most bacteria, we calculate the “essential metabolites” for each bacterium, identifying a total of 215 essential metabolites that primarily include ions, amino acids, nucleotides, vitamins, and sugars. “Essential metabolites” are defined as metabolites necessary for bacterial growth, thus necessitating their acquisition from the medium. Metabolites that are essential for more than 10% of the bacteria are added to the medium. We then label the carbon sources according to the Biolog phenotype array^78^ and limit their flux to reduce selective utilization. We also add the corresponding substrate to the medium. Finally, glucose is added when the substrate is not a carbon source. Depending on the different oxygen requirements, medium can be divided into four categories: with both oxygen and glucose, with oxygen only, with glucose only, and with neither oxygen nor glucose. All the medium used in this manuscript are available at GitHub (https://github.com/allons42/eBiota).

For bacteria of the same species with pathway similarity over 95% and differences in growth and maximal production efficiency less than 5%, only the one with highest production efficiency is retained. Of the 309 substrates, 191 carbon, nitrogen, phosphorus, and sulfur sources labeled by Biolog, as well as polysaccharides and fatty acids, were used for subsequent co-culture simulations. We use 83 intermediates, either essential metabolites or can be utilized and produced by more than 10% of the bacteria, excluding lipopolysaccharides, peptidoglycans, and inorganic compounds. The full results are shown in the **Supplementary Table S3** and on our website. ProdFBA was implemented in python 3.7 using COBRApy^79^ package 0.22.0 and the Gurobi solve (https://www.gurobi.com, Gurobi Optimizer v10.0.2). MICOM was performed by MICOM^9^ 0.37.0 with the Gurobi optimizer.

### The implementation of DeepCooc

We downloaded a total of 23,323 microbial samples from EMP (https://earthmicrobiome.org). We applied the HiOrCo method (https://github.com/cdanielmachado/hiorco)^30^ to capture co-occurring and mutually excluding members based on these microbial samples for training DeepCooc. Co-occurring (or mutually excluding) members were defined as bacteria that occur together across samples at a significantly higher (or lower) frequency than would be expected by chance. The resulting dataset was split into training, validation, and test sets at an 18:1:1 ratio. Since we used 16S rRNA data, DeepCooc currently operates at a genus-level resolution. While using metagenomic data could enhance taxonomic resolution, doing so would require a substantially larger sample size due to the exponential increase in taxa combination when moving from genus to strain level.

DeepCooc is invariant to the input order of bacteria. Specifically, we perform a full-permutation on the order of the input GEMs, and feed all full-permutation batched data into the DeepCooc. We minimize an auxiliary loss to minimize the differences among predictions across all permutations. For each permutation, the DeepCooc-single module receives the transport reaction trait matrix of each GEM as input. This matrix, representing 335 transport reactions found in eBiota-GEM that export metabolites to extracellular spaces, has rows representing these reactions and columns representing four traits: upper bound, lower bound, charge of the transported metabolite, and the number of reactions it participates in. We use transport reactions and charge of their cargos according to previous studies^80–82^, which demonstrate the efficacy of transport reactions in predicting community co-occurrence and how the charge of molecules influences their transportation. DeepCooc was implemented in Python 3.7, with its architecture detailed in **Extended Data Fig. 4**. We applied incremental learning to reduce the need to load the entire large dataset into memory simultaneously. The default training parameters for DeepCooc were as follows: Adam optimizer, binary cross-entropy (BCE) loss, learning rate of 0.0001, batch size of 128, 100 epochs, and an early stopping patience of 15 epochs. The parameter files of DeepCooc is available at Zenodo (https://zenodo.org/records/13895108). DeepCooc was compared with three commonly used machine learning methods: logistic regression, support vector machine, and naive Bayes. All these models were implemented using the scikit-learn Python library. Specifically, both logistic regression and support vector machine were implemented using the SGDClassifier function, with the former employing stochastic gradient descent (SGD) with a log loss function and the latter utilizing SGD with a modified Huber loss function. Naive Bayes was implemented as a Bernoulli Naive Bayes classifier using the BernoulliNB function. The random forest algorithm is not included because its default implementation in Python does not support incremental learning.

### The implementation of eBiota to various cases

To collect experimental data on production of various metabolites through microbial processes, we conducted a comprehensive search on PubMed and Google Scholar. If the GEMs for bacteria from these studies were not available in our eBiota-GEM database, we reconstructed the GEMs for each bacterium using either bacterial assemblies from the same species (or genus) download from RefSeq or by searching for the closest bacterial assemblies through the similarity of 16S rRNA sequence alignment. Based on the eBiota workflow, essential metabolites were added to the medium, which was further adjusted according to specific experimental conditions described in the study. We then used eBiota, applying CoreBFS to filter strains with complete pathways, ProdFBA to identify strains with high hydrogen production efficiency, and ProdFBA again to design high-efficiency communities. DeepCooc was then used to predict co-occurrence patterns for 2- and 3-member communities. Finally, we obtained community composition, production efficiency, cross-feeding, interactions, and co-occurrence. The accession number, GEMs, medium, substrates, designed communities, and results for these cases are publicly available in the Supplementary Tables, GitHub (https://github.com/allons42/eBiota), and Zenodo (https://zenodo.org/records/13895108).

### *In vitro* pairwise culture experiments

Tomato rhizosphere soils were sampled and sequenced in our previous studies^64^. One representative strain was selected from each species to minimize redundancy, yielding a final set of 94 bacterial strains together with *R. solanacearum* QL-Rs1115. The bacteria preserved in glycerol stocks were streaked onto Tryptic Soy Agar plates for activation, followed by 48-hour incubation to obtain single colonies. Single colonies were then transferred into 96-well plates containing 200 μL of Tryptic Soy Broth each and incubated on a shaker at 170 rpm and 30°C for 24 hours. The bacterial suspensions were adjusted to an OD_600_ of 0.5 using sterile water.

Subsequently, 2 μL of each bacterial suspension was transferred into 96-well plates containing Nutrient Agar (NA) medium and incubated at 30°C and 170 rpm for 48 hours. Fermentation broth was obtained by centrifuging the 96-well plate. Next, 20 μL of fermentation broth (with sterile water as control) was transferred into 96-well plates containing 178 μL of 10% concentration NA liquid medium. Subsequently, 2 μL of *R. solanacearum* QL-Rs1115 bacterial suspension (OD_600_ = 1.0) was inoculated into each well, followed by incubation at 30°C and 170 rpm for 24 hours. The *R. solanacearum* QL-Rs1115 density at 600 nm wavelength was measured using a microplate reader. The competition intensity of the bacterial strains against *R. solanacearum* was expressed as [- (OD_600_sequential-culture_ - OD_600_mono-culture_) / OD_600_mono-culture_]. OD_600_sequential-culture_ and OD_600_mono-culture_ represent the absorbance at OD600 in the sequential cultures or mono-culture, respectively.

### *In vitro* co-culture experiments of 94-strains bacterial communities

After incubation, equal volumes of bacterial suspensions from each of 94 strains were mixed to create a synthetic bacterial community suspension. This synthetic community suspension was inoculated at 1% volume into 96-well plates with LB medium. The plates were then incubated on a shaker at 170 rpm and 30°C. At time points 0 hours, 12 hours, 24 hours, 36 hours, and 48 hours, 1 mL of bacterial culture was sampled from each well and transferred to 1 mL sterile centrifuge tubes, followed by immediate freezing in liquid nitrogen. The samples were then sent to Shanghai Biozeron Biological Technology Co. Ltd. (Shanghai, China) for 16S rRNA gene sequencing. Based on the sequencing results, the growth dynamics of individual bacterial species within the community were analyzed.

### 16S rRNA gene sequencing and analysis

Microbial DNA was extracted from 25 samples using the E.Z.N.A.® Soil DNA Kit (Omega Bio-tek, Norcross, GA, U.S.) according to manufacturer’s protocols. The V4-V5 region of the bacteria 16S ribosomal RNA gene were amplified by PCR (95 °C for 2 min, followed by 25 cycles at 95 °C for 30 s, 55 °C for 30 s, and 72 °C for 30 s and a final extension at 72 °C for 5 min) using primers 515F (5’-barcode-GTGCCAGCMGCCGCGG-3’) and 907R (5’-CCGTCAATTCMTTTRAGTTT-3’), where barcode is an eight-base sequence unique to each sample. PCR reactions were performed in triplicate 20 μL mixture containing 4 μL of 5 × FastPfu Buffer, 2 μL of 2.5 mM dNTPs, 0.8 μL of each primer (5 μM), 0.4 μL of FastPfu Polymerase, and 10 ng of template DNA. Amplicons were extracted from 2% agarose gels and purified using the AxyPrep DNA Gel Extraction Kit (Axygen Biosciences, Union City, CA, U.S.) according to the manufacturer’s instructions.

Purified PCR products were quantified by Qubit®3.0 (Life Invitrogen) and every twenty-four amplicons whose barcodes were different were mixed equally. The pooled DNA product was used to construct Illumina Pair-End library following Illumina’s genomic DNA library preparation procedure. Then the amplicon library was paired-end sequenced (2 × 250) on an Illumina MiSeq platform (Shanghai BIOZERON Co., Ltd) according to the standard protocols.

Raw fastq files were first demultiplexed using Perl scripts according to the barcode sequences information for each sample with the following criteria: (i) The 250 bp reads were truncated at any site receiving an average quality score < 20 over a 10 bp sliding window, discarding the truncated reads that were shorter than 50 bp. (ii) exact barcode matching, 2 nucleotide mismatches in primer matching, reads containing ambiguous characters were removed. (iii) only sequences that overlap longer than 10 bp were assembled according to their overlap sequence. Reads which could not be assembled were discarded. OTUs were clustered with 97% similarity cutoff using UPARSE (version 7.1 http://drive5.com/uparse/) and chimeric sequences were identified and removed using UCHIME. The phylogenetic affiliation of each 16S rRNA gene sequence was analyzed by RDP Classifier (http://rdp.cme.msu.edu/) against the SILVA (SSU132) 16S rRNA database using confidence threshold of 70% or uclust algorithm (http://www.drive5.com/usearch/manual/uclust_algo.html) against the SILVA (SSU138.1) 16S rRNA database using the confidence threshold of 80%.

### The implementation of eBiota to *in vitro* co-cultures

We analyzed the growth (measured as OD_600_) of the synthetic bacterial community comprising 94 strains and found that maximum growth was achieved at 12 hours. Based on the definition of FBA, we used eBiota to simulate this community in LB medium and calculated the cumulative relative abundance from 0 to 12 hours. We compared these cumulative results with the experimental relative abundance data obtained at 12 hours.

### Pareto front analysis

In multi-objective optimization, the Pareto front refers to the set of solutions that provide optimal trade-offs across all objective functions and is widely applicable in fields such as economics, engineering, and manufacturing. A solution lies on the Pareto front if it is not dominated by any other solution in the feasible space. In the simulated co-culture of a strain with *R. solanacearum* QL-Rs1115, each solution corresponds to the maximum growth of *R. solanacearum* QL-Rs1115 when the growth of the co-cultured strain is constrained to be no less than a defined threshold. This process was repeated 24 times under different thresholds. The line fitted to these solutions (points) represents the Pareto front, illustrating the balance between the growth of the two strains. Therefore, A Pareto front displaying a negative linear trend (*e.g.*, CUR1, ART3, BAC1, BUR1, FIC1, STA1, EMP1, and FLA5 in **Fig. 6B**) indicates a competitive relationship between the two strains. Conversely, a Pareto front approximating a right angle (*e.g.*, FLA5, EMP1, CHR3 and ESC1 in **Fig. 6B**) suggests a weak association, or even potential independence, between the two strains.

### Statistical analysis

All statistical analyses and plots were implements using R (version 4.0.3). Paired and unpaired continuous variables, were represented by mean ± standard deviation and tested by the two-tailed Wilcoxon signed-rank and rank-sum tests, respectively. Statistical significance was considered when *p* value < 0.05. PERMANOVA was used to test the significance differences in the profiles among groups. PhyloMint (version 0.1.0)^68^ was used to calculate metabolic competition index, complementarity index, and a derived metabolic distance score between all strains and *R. solanacearum* QL-Rs1115.

## Data availability

Accessions for publicly available genomic data are given in the Supplementary Tables. More results and media are available at GitHub (https://github.com/allons42/eBiota) and Zenodo (https://zenodo.org/records/13895108). Sequencing data of 94 bacterial strains and *R. solanacearum* QL-Rs1115 have been deposited in Zenodo (https://zenodo.org/records/13764123). Raw sequencing data of the 25 *in vitro* culture samples are available at the National Center for Biotechnology Information database (accession number: PRJNA1163256).

## Code availability

The code for eBiota with CoreBFS, ProdFBA, and DeepCooc in this manuscript is available at https://github.com/allons42/eBiota.

## Author contributions

HQZ, XQJ and ZW co-supervised the study. XQJ, HQZ, JHH and HYZ designed the study. XQJ, JHH, HYZ, JYG, and YLL carried out the eBiota platform and algorithms. SHG, XRY and ZW conducted the experiments. XQJ, JHH, HYZ, SHG, XRY, PXG, YYZ, QG, CHW and ML ran the analyses and plotted the figures. XQJ, JHH, HYZ, SHG, AJ and HQZ wrote and revised the manuscript. All the authors read and accepted the final version of the manuscript.

## Acknowledgments

This work was supported by the National Science and Technology Major Project (2025ZD01901200, 2025ZD01901800) and the National Natural Science Foundation of China (32570752, 32300078, 42325704). Part of the analysis was performed on the High-Performance Computing Platform of Peking University, and Biomedical Computing Platform of National Biomedical Imaging Center of Peking University.

## References

1. Yan, W., Cao, Z., Ding, M. & Yuan, Y. Design and construction of microbial cell factories based on systems biology. Synth. Syst. Biotechnol. 8, 176–185 (2023).

2. van Leeuwen, P. T., Brul, S., Zhang, J. & Wortel, M. T. Synthetic microbial communities (SynComs) of the human gut: design, assembly, and applications. FEMS Microbiol. Rev. 47, fuad012 (2023).

3. Gelsinger, D. R. et al. Metagenomic editing of commensal bacteria in vivo using CRISPR-associated transposases. Science 390, eadx7604 (2025).

4. Qin, L., Liu, X., Xu, K. & Li, C. Mining and design of biosensors for engineering microbial cell factory. Curr. Opin. Biotech. 75, 102694 (2022).

5. Madhavan, A. et al. Design and genome engineering of microbial cell factories for efficient conversion of lignocellulose to fuel. Bioresour. Technol. 370, 128555 (2023).

6. Ergal, İ., et al. Biohydrogen production beyond the Thauer limit by precision design of artificial microbial consortia. Commun. Biol. 3, 1–12 (2020).

7. García-Jiménez, B., Torres-Bacete, J. & Nogales, J. Metabolic modelling approaches for describing and engineering microbial communities. Comput. Struct. Biotechnol. J. 19, 226–246 (2021).

8. Liu, Y.-Y. Controlling the human microbiome. Cell Syst. 14, 135–159 (2023).

9. Diener, C., Gibbons, S. M. & Resendis-Antonio, O. MICOM: Metagenome-Scale Modeling To Infer Metabolic Interactions in the Gut Microbiota. mSystems 5, e00606–19 (2020).

10. Zomorrodi, A. R. & Maranas, C. D. OptCom: a multi-level optimization framework for the metabolic modeling and analysis of microbial communities. PLoS Comput. Biol. 8, e1002363 (2012).

11. Chan, S. H. J., Simons, M. N. & Maranas, C. D. SteadyCom: Predicting microbial abundances while ensuring community stability. PLoS Comput. Biol. 13, e1005539 (2017).

12. Baldini, F. et al. The Microbiome Modeling Toolbox: from microbial interactions to personalized microbial communities. Lect. Notes. Comput. Sc. 35, 2332–2334 (2019).

13. Li, P., Roos, S., Luo, H., Ji, B. & Nielsen, J. Metabolic engineering of human gut microbiome: Recent developments and future perspectives. Metab. Eng. 79, 1–13 (2023).

14. Marcelino, V. R. et al. Disease-specific loss of microbial cross-feeding interactions in the human gut. Nat. Commun. 14, 6546 (2023).

15. Quinn-Bohmann, N. et al. Microbial community-scale metabolic modelling predicts personalized short-chain fatty acid production profiles in the human gut. Nat. Microbiol. 9, 1700–1712 (2024).

16. Ruan, Z. et al. Engineering natural microbiomes toward enhanced bioremediation by microbiome modeling. Nat. Commun. 15, 4694 (2024).

17. Eng, A. & Borenstein, E. An algorithm for designing minimal microbial communities with desired metabolic capacities. Bioinformatics 32, 2008–2016 (2016).

18. Julien-Laferrière, A. et al. A Combinatorial Algorithm for Microbial Consortia Synthetic Design. Sci. Rep. 6, 29182 (2016).

19. García-Jiménez, B., García, J. L. & Nogales, J. FLYCOP: metabolic modeling-based analysis and engineering microbial communities. Bioinformatics 34, i954–i963 (2018).

20. Eng, A. & Borenstein, E. Microbial community design: methods, applications, and opportunities. Curr. Opin. Biotech. 58, 117–128 (2019).

21. Orth, J. D., Thiele, I. & Palsson, B. Ø. What is flux balance analysis? Nat. Biotechnol. 28, 245–248 (2010).

22. O’Brien, E. J., Monk, J. M. & Palsson, B. O. Using Genome-scale Models to Predict Biological Capabilities. Cell 161, 971–987 (2015).

23. Lewis, N. E. et al. Omic data from evolved E. coli are consistent with computed optimal growth from genome-scale models. Mol. Syst. Biol. 6, 390 (2010).

24. Faure, L., Mollet, B., Liebermeister, W. & Faulon, J.-L. A neural-mechanistic hybrid approach improving the predictive power of genome-scale metabolic models. Nat. Commun. 14, 4669 (2023).

25. Klamt, S. & Mahadevan, R. On the feasibility of growth-coupled product synthesis in microbial strains. Metab. Eng. 30, 166–178 (2015).

26. Kenefake, D., Armingol, E., Lewis, N. E. & Pistikopoulos, E. N. An improved algorithm for flux variability analysis. BMC Bioinformatics 23, 550 (2022).

27. Burgard, A. P., Vaidyaraman, S. & Maranas, C. D. Minimal reaction sets for Escherichia coli metabolism under different growth requirements and uptake environments. Biotechnol. Prog. 17, 791–797 (2001).

28. Wang, M. et al. Even allocation of benefits stabilizes microbial community engaged in metabolic division of labor. Cell Rep. 40, 111410 (2022).

29. Jiang, X. et al. How Microbes Shape Their Communities? A Microbial Community Model Based on Functional Genes. Genomics Proteomics Bioinformatics 17, 91–105 (2019).

30. Machado, D. et al. Polarization of microbial communities between competitive and cooperative metabolism. Nat. Ecol. Evol. 5, 195–203 (2021).

31. Li, D. et al. Coexistence patterns of soil methanogens are closely tied to methane generation and community assembly in rice paddies. Microbiome 9, 20 (2021).

32. Eiler, A., Heinrich, F. & Bertilsson, S. Coherent dynamics and association networks among lake bacterioplankton taxa. ISME J. 6, 330–342 (2012).

33. Kurtz, Z. D. et al. Sparse and compositionally robust inference of microbial ecological networks. PLoS Comput. Biol. 11, e1004226 (2015).

34. Friedman, J. & Alm, E. J. Inferring correlation networks from genomic survey data. PLoS Comput. Biol. 8, e1002687 (2012).

35. Kamneva, O. K. Genome composition and phylogeny of microbes predict their co-occurrence in the environment. PLoS Comput. Biol. 13, e1005366 (2017).

36. Cerini, F., Vignoli, L., Blust, M. & Strona, G. Functional traits predict species co-occurrence patterns in a North American Odonata metacommunity. Ecosphere 14, e4732 (2023).

37. Lieven, C. et al. MEMOTE for standardized genome-scale metabolic model testing. Nat. Biotechnol. 38, 272–276 (2020).

38. Magnúsdóttir, S. et al. Generation of genome-scale metabolic reconstructions for 773 members of the human gut microbiota. Nat. Biotechnol. 35, 81–89 (2017).

39. King, Z. A. et al. BiGG Models: A platform for integrating, standardizing and sharing genome-scale models. Nucleic Acids Res. 44, D515–D522 (2016).

40. Mahadevan, R. & Palsson, B. O. Properties of metabolic networks: structure versus function. Biophys. J. 88, L07–09 (2005).

41. Broido, A. D. & Clauset, A. Scale-free networks are rare. Nat. Commun. 10, 1017 (2019).

42. Albert, R. Scale-free networks in cell biology. J. Cell Sci. 118, 4947–4957 (2005).

43. Schuchmann, K. & Müller, V. Autotrophy at the thermodynamic limit of life: a model for energy conservation in acetogenic bacteria. Nat. Rev. Microbiol. 12, 809–821 (2014).

44. Jing, E. et al. ECG Heartbeat Classification Based on an Improved ResNet-18 Model. Comput. Math Methods Med. 2021, 6649970 (2021).

45. Ge, F. et al. Prediction of disease-associated nsSNPs by integrating multi-scale ResNet models with deep feature fusion. Brief. Bioinform. 23, bbab530 (2022).

46. He, K., Zhang, X., Ren, S. & Sun, J. Deep Residual Learning for Image Recognition. in 2016 IEEE Conference on Computer Vision and Pattern Recognition (CVPR) 770-778 (IEEE, 2016).

47. Thompson, L. R. et al. A communal catalogue reveals Earth’s multiscale microbial diversity. Nature 551, 457–463 (2017).

48. Chen, C., Liao, C. & Liu, Y.-Y. Teasing out missing reactions in genome-scale metabolic networks through hypergraph learning. Nat. Commun. 14, 2375 (2023).

49. Djemai, K., Drancourt, M. & Tidjani Alou, M. Bacteria and Methanogens in the Human Microbiome: a Review of Syntrophic Interactions. Microb. Ecol. 83, 536–554 (2022).

50. Scott, W. T. et al. A structured evaluation of genome-scale constraint-based modeling tools for microbial consortia. PLoS Comput. Biol. 19, e1011363 (2023).

51. Podosokorskaya, O. A. et al. Tenuifilum thalassicum gen. nov., sp. nov., a novel moderate thermophilic anaerobic bacterium from a Kunashir Island shallow hot spring representing a new family Tenuifilaceae fam. nov. in the class Bacteroidia. Syst. Appl. Microbiol. 43, 126126 (2020).

52. Zeidan, A. A. & van Niel, E. W. J. A quantitative analysis of hydrogen production efficiency of the extreme thermophile *Caldicellulosiruptor owensensis* OLT. Int. J. Hydrog. Energy 35, 1128–1137 (2010).

53. Du, Y., Zou, W., Zhang, K., Ye, G. & Yang, J. Advances and Applications of Clostridium Co-culture Systems in Biotechnology. Front. Microbiol. 11, 560223 (2020).

54. Cui, Y., Yang, K.-L. & Zhou, K. Using Co-Culture to Functionalize Clostridium Fermentation. Trends Biotechnol. 39, 914–926 (2021).

55. Li, P. & Zhu, M. A consolidated bio-processing of ethanol from cassava pulp accompanied by hydrogen production. Bioresour. Technol. 102, 10471–10479 (2011).

56. Weetall, H. H., Sharma, B. P. & Detar, C. C. Photometabolic production of hydrogen from organic substrates by free and immobilized mixed cultures of Rhodospirillum rubrum and Klebsiella pneumoniae. Biotechnology and Bioengineering 23, 605–614 (1981).

57. Shi, X.-C. et al. Impact of electron scavenging during electric current generation from propionate by a *Geobacter* co-culture. Chemical Engineering Journal 418, 129357 (2021).

58. Masset, J. et al. Fermentative hydrogen production from glucose and starch using pure strains and artificial co-cultures ofClostridium spp. Biotechnol. Biofuels 5, 35 (2012).

59. Patel, S. K. S., Singh, M., Kumar, P., Purohit, H. J. & Kalia, V. C. Exploitation of defined bacterial cultures for production of hydrogen and polyhydroxybutyrate from pea-shells. Biomass Bioenerg. 36, 218–225 (2012).

60. Medlock, G. L. et al. Inferring Metabolic Mechanisms of Interaction within a Defined Gut Microbiota. Cell Syst. 7, 245–257.e7 (2018).

61. Wymore Brand, M., et al. The Altered Schaedler Flora: Continued Applications of a Defined Murine Microbial Community. ILAR J. 56, 169–178 (2015).

62. Kwak, M.-J. et al. Rhizosphere microbiome structure alters to enable wilt resistance in tomato. Nat. Biotechnol. 36, 1100–1109 (2018).

63. Compant, S., Samad, A., Faist, H. & Sessitsch, A. A review on the plant microbiome: Ecology, functions, and emerging trends in microbial application. J. Adv. Res. 19, 29–37 (2019).

64. Gu, S. et al. Competition for iron drives phytopathogen control by natural rhizosphere microbiomes. Nat. Microbiol. 5, 1002–1010 (2020).

65. Heinken, A. & Thiele, I. Anoxic Conditions Promote Species-Specific Mutualism between Gut Microbes In Silico. Appl. Environ. Microb. 81, 4049–4061 (2015).

66. Ibrahim, M. & Raman, K. Two-species community design of lactic acid bacteria for optimal production of lactate. Comput. Struct. Biotechnol. J. 19, 6039–6049 (2021).

67. Yin, Q. et al. Ecological dynamics of Enterobacteriaceae in the human gut microbiome across global populations. Nat. Microbiol. 10, 541–553 (2025).

68. Lam, T. J., Stamboulian, M., Han, W. & Ye, Y. Model-based and phylogenetically adjusted quantification of metabolic interaction between microbial species. PLoS Comput. Biol. 16, e1007951 (2020).

69. Hu, H. et al. Guided by the principles of microbiome engineering: Accomplishments and perspectives for environmental use. mLife 1, 382–398 (2022).

70. Machado, D., Andrejev, S., Tramontano, M. & Patil, K. R. Fast automated reconstruction of genome-scale metabolic models for microbial species and communities. Nucleic Acids Res. 46, 7542–7553 (2018).

71. Edgar, R. C. MUSCLE: multiple sequence alignment with high accuracy and high throughput. Nucleic Acids Res 32, 1792–1797 (2004).

72. Price, M. N., Dehal, P. S. & Arkin, A. P. FastTree: computing large minimum evolution trees with profiles instead of a distance matrix. Mol Biol Evol 26, 1641–1650 (2009).

73. Reimer, L. C. et al. BacDive in 2022: the knowledge base for standardized bacterial and archaeal data. Nucleic Acids Res. 50, D741–D746 (2022).

74. Gerlee, P., Lizana, L. & Sneppen, K. Pathway identification by network pruning in the metabolic network of Escherichia coli. Bioinformatics 25, 3282–3288 (2009).

75. Wagner, A. & Fell, D. A. The small world inside large metabolic networks. Proc. Biol. Sci. 268, 1803–1810 (2001).

76. Ravikrishnan, A., Nasre, M. & Raman, K. Enumerating all possible biosynthetic pathways in metabolic networks. Sci. Rep. 8, 9932 (2018).

77. Gebser, M., Kaminski, R., Kaufmann, B. & Schaub, T. Clingo = ASP + Control: Preliminary Report. in Technical Communications of the Thirtieth International Conference on Logic Programming (ICLP’14) 1-9 (ICLP, 2014).

78. Bochner, B. R., Gadzinski, P. & Panomitros, E. Phenotype MicroArrays for High-Throughput Phenotypic Testing and Assay of Gene Function. Genome Res. 11, 1246–1255 (2001).

79. Ebrahim, A., Lerman, J. A., Palsson, B. O. & Hyduke, D. R. COBRApy: COnstraints-Based Reconstruction and Analysis for Python. BMC Syst. Biol 7, 74 (2013).

80. DiMucci, D., Kon, M. & Segrè, D. Machine Learning Reveals Missing Edges and Putative Interaction Mechanisms in Microbial Ecosystem Networks. mSystems 3, e00181–18 (2018).

81. Erkus, O. et al. Multifactorial diversity sustains microbial community stability. ISME J. 7, 2126–2136 (2013).

82. D’Souza, G. et al. Ecology and evolution of metabolic cross-feeding interactions in bacteria. Nat. Prod. Rep. 35, 455–488 (2018).

