## Supplementary for "eBiota: Designing microbial communities from large seed pools with desired function using rapid optimization and deep learning"

Extended Data Figures

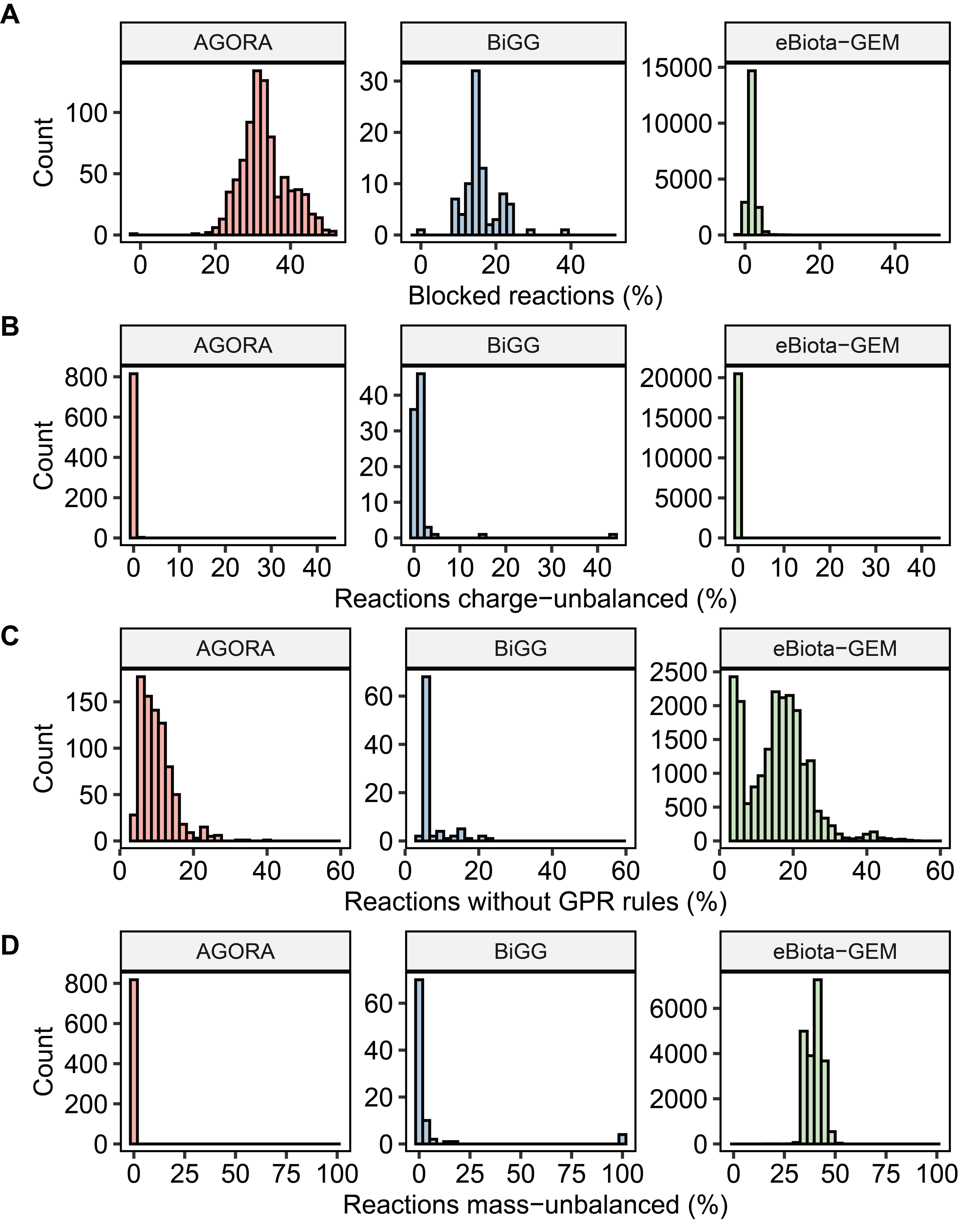

Extended Data Fig. 1 The quality of GEMs in eBiota-GEM. Histogram plots show the distribution of reactions blocked (A), charge-unbalanced (B), without gene-protein-reaction (GPR) rules (C) and mass-unbalanced (D) among GEMs in the AGORA, BiGG and eBiota-GEM. GEMs in the eBiota-GEM have fewer of blocked reactions and charge-unbalanced reactions, with slightly inferior performance in terms of reactions with GPR rules and mass-balanced, compared to AGORA and BiGG.

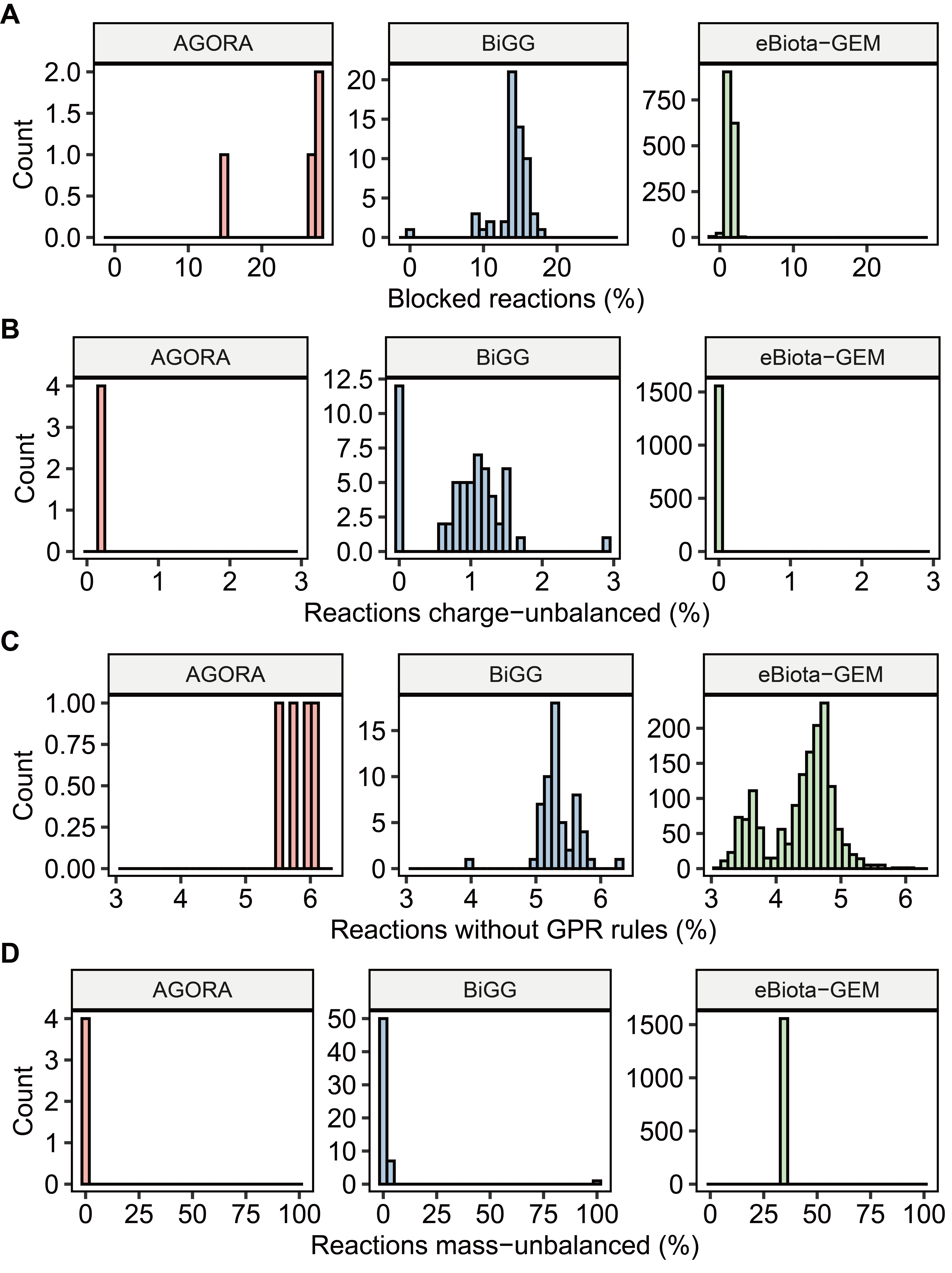

Extended Data Fig. 2 The quality of *E*. *coli* GEMs in eBiota-GEM. Histogram plots show the distribution of reactions blocked (A), charge-unbalanced (B), without GPR rules (C) and mass-unbalanced (D) among the *E. coli* GEMs within the AGORA, BiGG and eBiota-GEM. To compare these resources at the species level, we selected *Escherichia coli*, the only species with more than three GEMs in each resource (4 in AGORA, 58 in BiGG, and 1,557 in eBiota-GEM). GEMs in eBiota-GEM have fewer blocked reactions, charge-unbalanced reactions, and reactions without GPR rules, but more mass-unbalanced reactions compared to AGORA and BiGG.

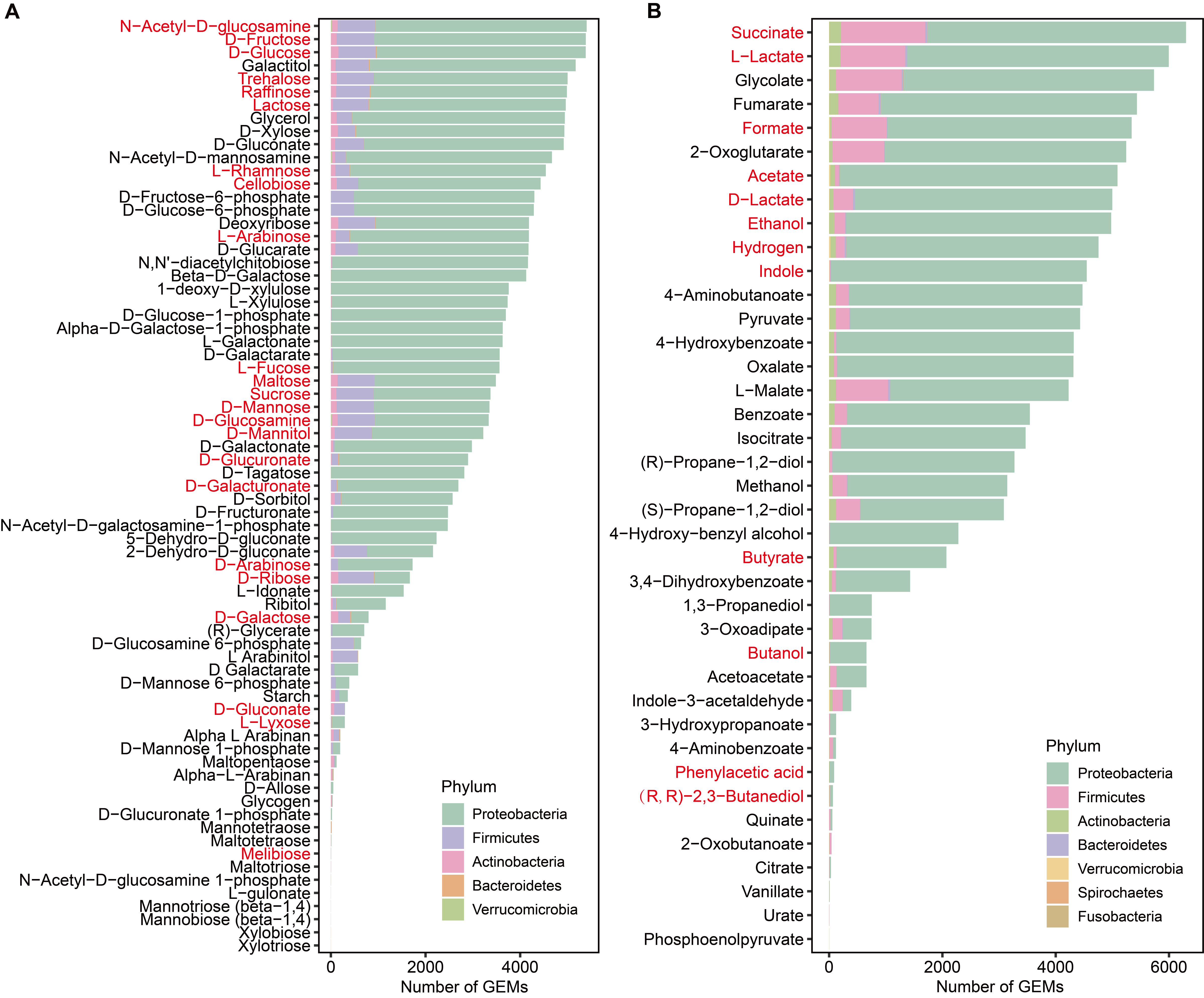

Extended Data Fig. 3 Results of ProdFBA. Bar plots show the number of gut bacteria (GEM) in each phylum that can consume the simple sugars (A) or secrete the fermentation products (B). Red: simple sugars or fermentation products that have been reported in the simulations of 773 gut bacteria.

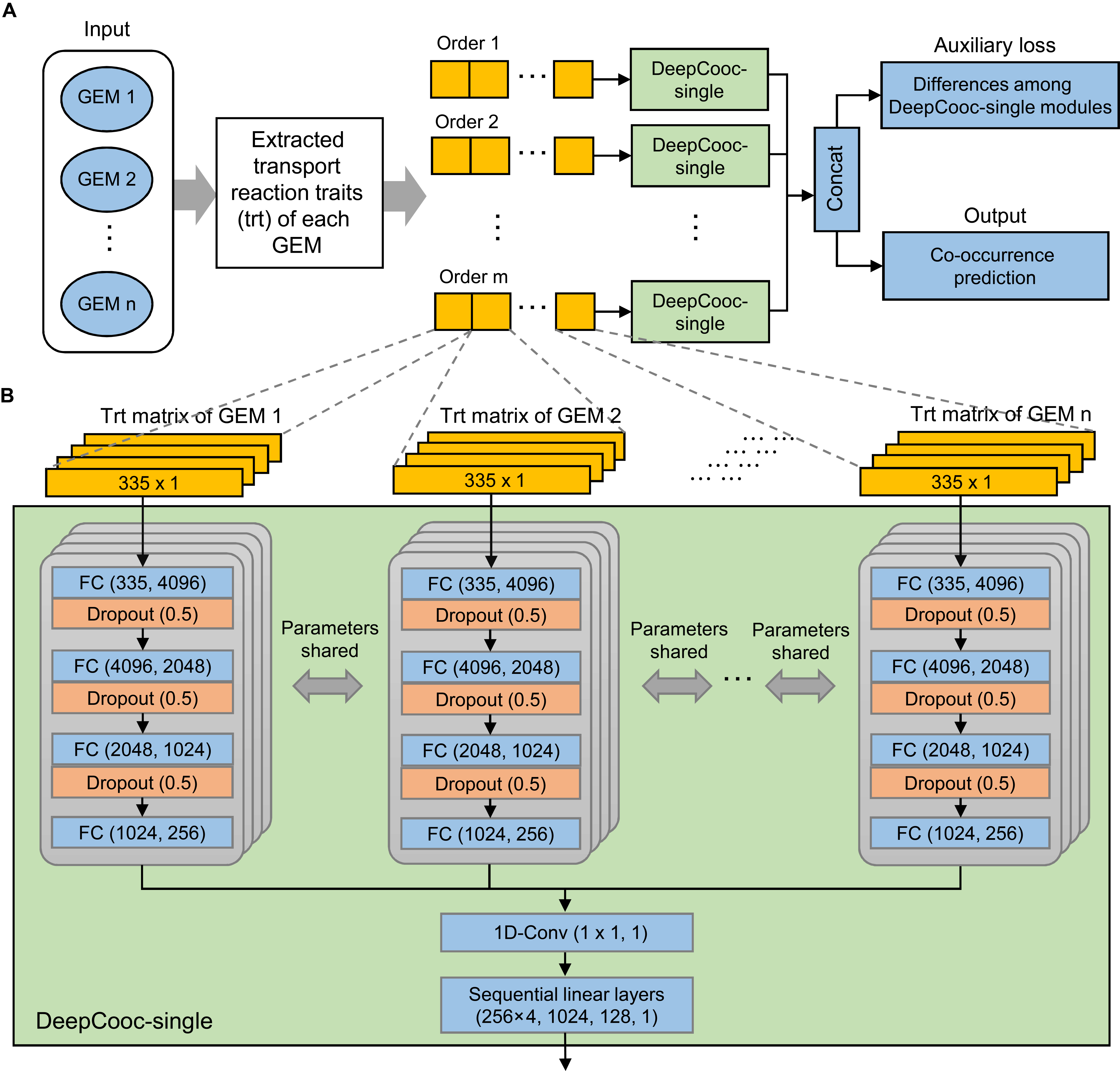

**Extended Data Fig. 4 Architecture of DeepCooc.** (**A**) The main architecture of DeepCooc. (**B**) The details of DeepCooc-single module. The transport reactions traits matrix composed of four traits of each of 335 transport reactions. These four traits of transport reactions include upper bound, lower bound, charge of the transported metabolite and the number of reactions it participates in. Each sequential linear layer was followed by a dropout layer (ratio = 0.5) and leaky rectified linear unit (ReLU, alpha = 0.1). FC: A fully connected layer. 1D-Conv: A 1-D convolutional layer.

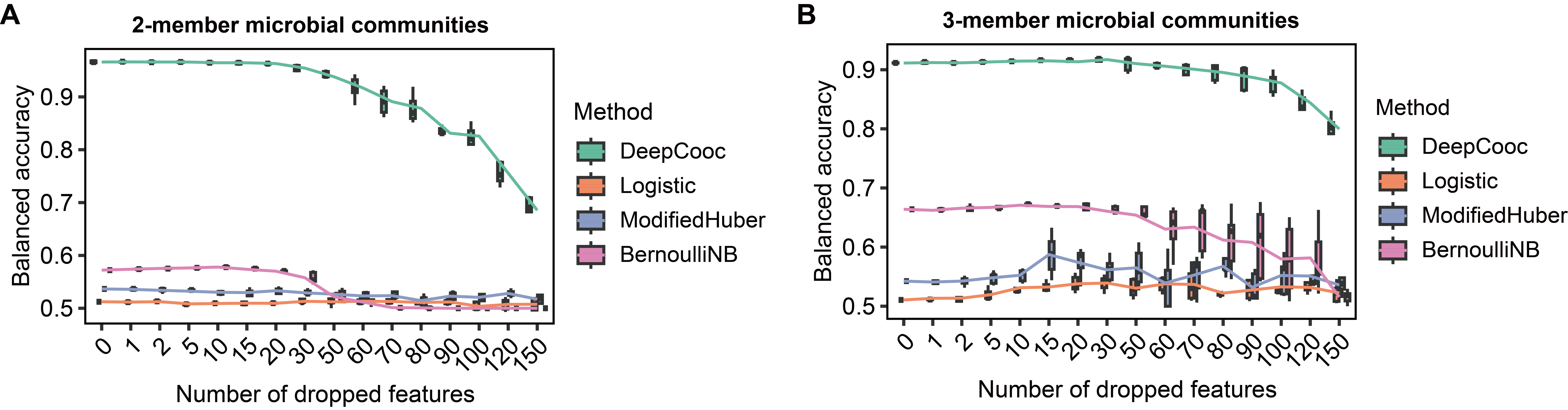
**Extended Data Fig. 5 Comparison of performance of DeepCooc with other methods.** Changes in the AUROC scores of four methods when varying the number of features dropped, applied to microbial communities consisting of two (**A**) and three (**B**) members. The total number of features is 335. logistic: logistic regression; SVM: support vector machine. Naive Bayes: Bernoulli naive bayes.

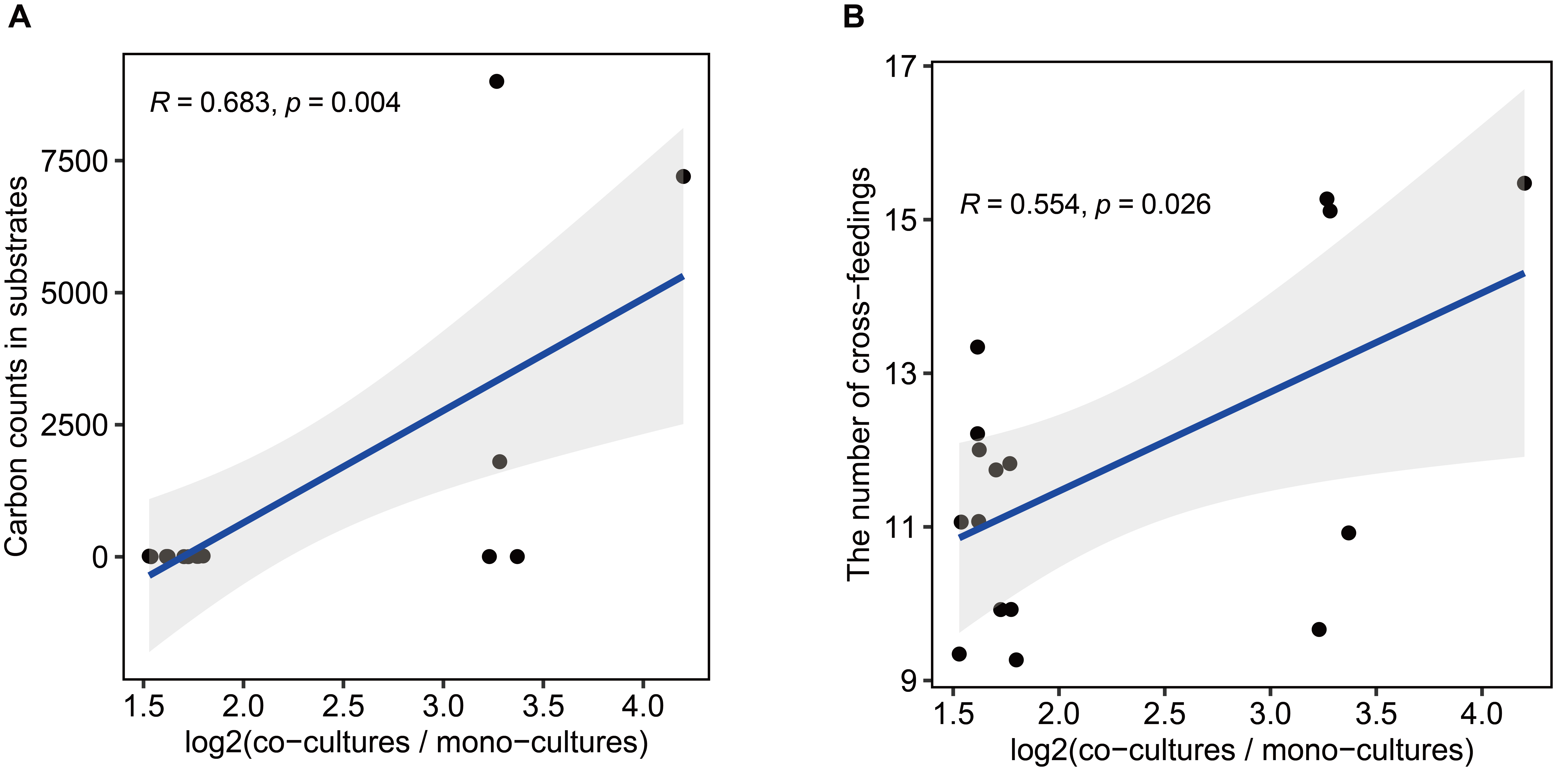

**Extended Data Fig. 6 Relationship of fold change in hydrogen production efficiency with substrates and cross-feedings.** The fold change in hydrogen production efficiency between co-cultures and mono-cultures is positively correlated with the number of carbon atoms in the substrates (**A**) and the number of cross-feeding substances (**B**). Pearson correlation coefficient, *p* value and a linear regression line with 95% confidence intervals are shown.

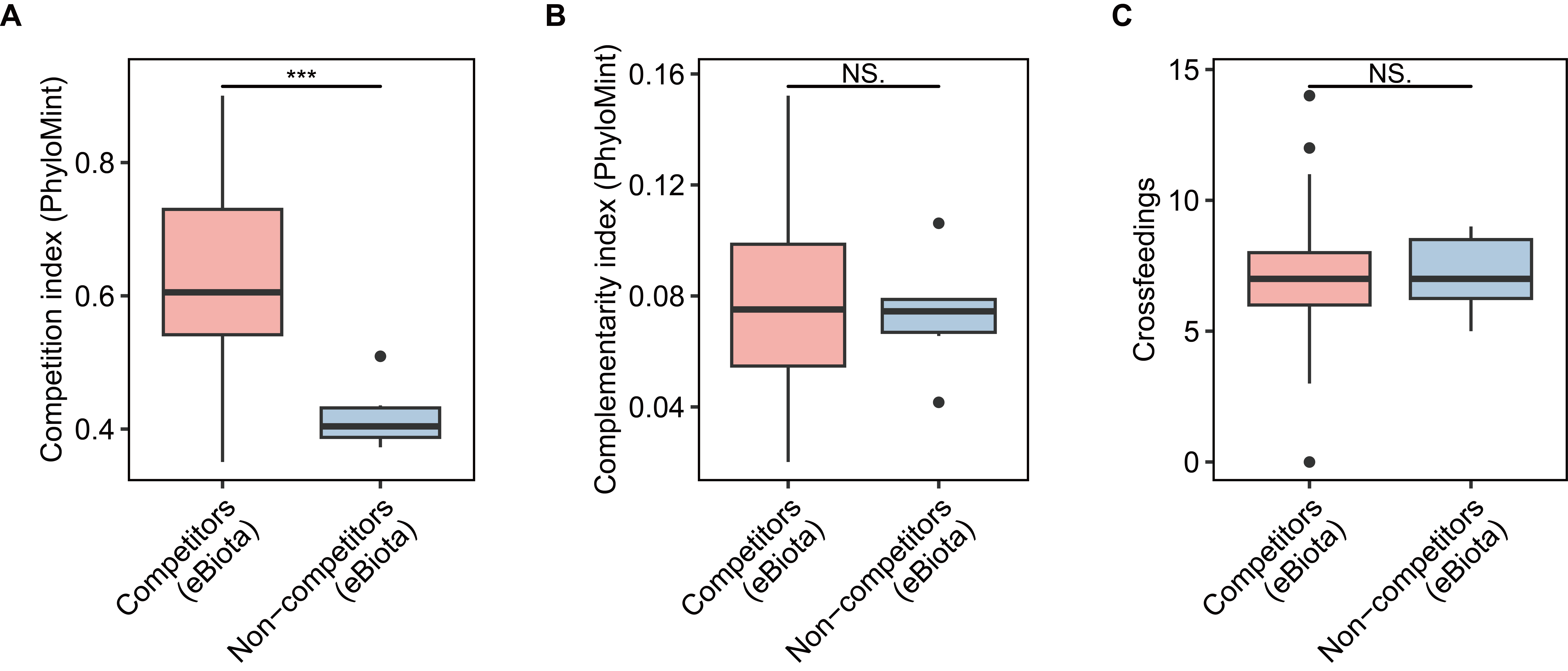

**Extended Data Fig. 7** **Comparison of metabolic** **characteristics of competitors and non-competitors.** Comparison of the competition index (**A**) and complementary index (**B**) for competitors and non-competitors. These indices were calculated using PhyloMint. (**C**) Comparison of the number of crossfeedings for these two groups. Competitors and non-competitors were identified using eBiota (competitors: n = 88; non-competitors: n = 6). ****p* < 0.001, Wilcoxon rank-sum test. NS: not significant.

In addition, there are nine Supplementary Table Files:

**Supplementary Table S1** Information on 21,514 bacteria in the eBiota-GEM.

**Supplementary Table S2** Pathway completeness for hydrogen production calculated by CoreBFS and MultiPus, evaluated using ProdFBA.

**Supplementary Table S3** Information on the simulated cultures of 21,514 bacteria using eBiota.

**Supplementary Table S4** Hydrogen production efficiency of *Tenuifilum thalassicum* 38H-str calculated by eBiota.

**Supplementary Table S5** Hydrogen production capabilities calculated by eBiota for pairwise co-cultures of taxa.

**Supplementary Table S6** Hydrogen production capabilities calculated by eBiota for co-culture of three taxa.

**Supplementary Table S7** Hydrogen yield calculated by eBiota for co-culture of six taxa selected from a 11-taxa pool.

**Supplementary Table S8** Production capabilities of 28 metabolites in microbial communities calculated with MICOM.

**Supplementary Table S9** Information on 94 bacterial strains isolated from rhizosphere and the plant pathogen *R. solanacearum* QL-Rs1115.
